# Platelet-endothelial crosstalk drives distinct endothelial dysfunction in bicuspid aortic valve disease

**DOI:** 10.64898/2026.11.04.763940

**Authors:** Luisa Weiss, Sarvesh Chelvanambi, Amarri Harrison, Taku Kasai, Yuto Nakamura, Mandy E Turner, Mark C Blaser, Marie Billaud, Masanori Aikawa, Sasha A Singh, Elena Aikawa

## Abstract

**Background:** Bicuspid aortic valves (BAV) predispose individuals to premature development of aortic stenosis (AS). The abnormal hemodynamic environment of BAV affects both platelets and valvular endothelial cells (VECs), but whether platelet-derived signals contribute to valve morphology-specific endothelial dysfunction remains unknown.

**Methods:** Platelets and VECs were obtained from patients with BAVs or tricuspid aortic valves (TAVs) undergoing valve replacement for AS. Platelets were activated to generate platelet releasate (PR). PR composition and its effects on VECs were investigated using complementary molecular, multi-omic, functional, and computational approaches.

**Results:** BAV and TAV patients exhibited markedly distinct PR profiles, which translated into differential endothelial responses. Strikingly, BAV-derived PR elicited a substantially more pronounced endothelial dysfunction phenotype, characterized by widespread molecular reprogramming and functional alterations. Integrated multi-omics uncovered mitochondrial dysfunction as a central feature of BAV PR-induced endothelial injury, accompanied by activation of pathways spanning endothelial remodeling, migration, and immune-cell recruitment. These molecular signatures translated into mitochondrial depolarization, altered mitochondrial dynamics, enhanced VEC migration, and increased neutrophil adhesion and chemotaxis. Finally, computational ligand-receptor and ligand-target analyses identified candidate signaling axes linking platelet-derived mediators to endothelial reprogramming, highlighting potential therapeutic targets.

**Conclusions:** We provide the first direct evidence that platelet activation contributes to AS initiation through the induction of endothelial dysfunction and that these effects are amplified in BAV. These findings identify platelet-VEC crosstalk as a previously unrecognized mechanism linking abnormal valve hemodynamics to disease onset. Targeting platelet-derived mediators may represent a novel therapeutic strategy for AS, specifically in BAV.

**Graphical Abstract:** 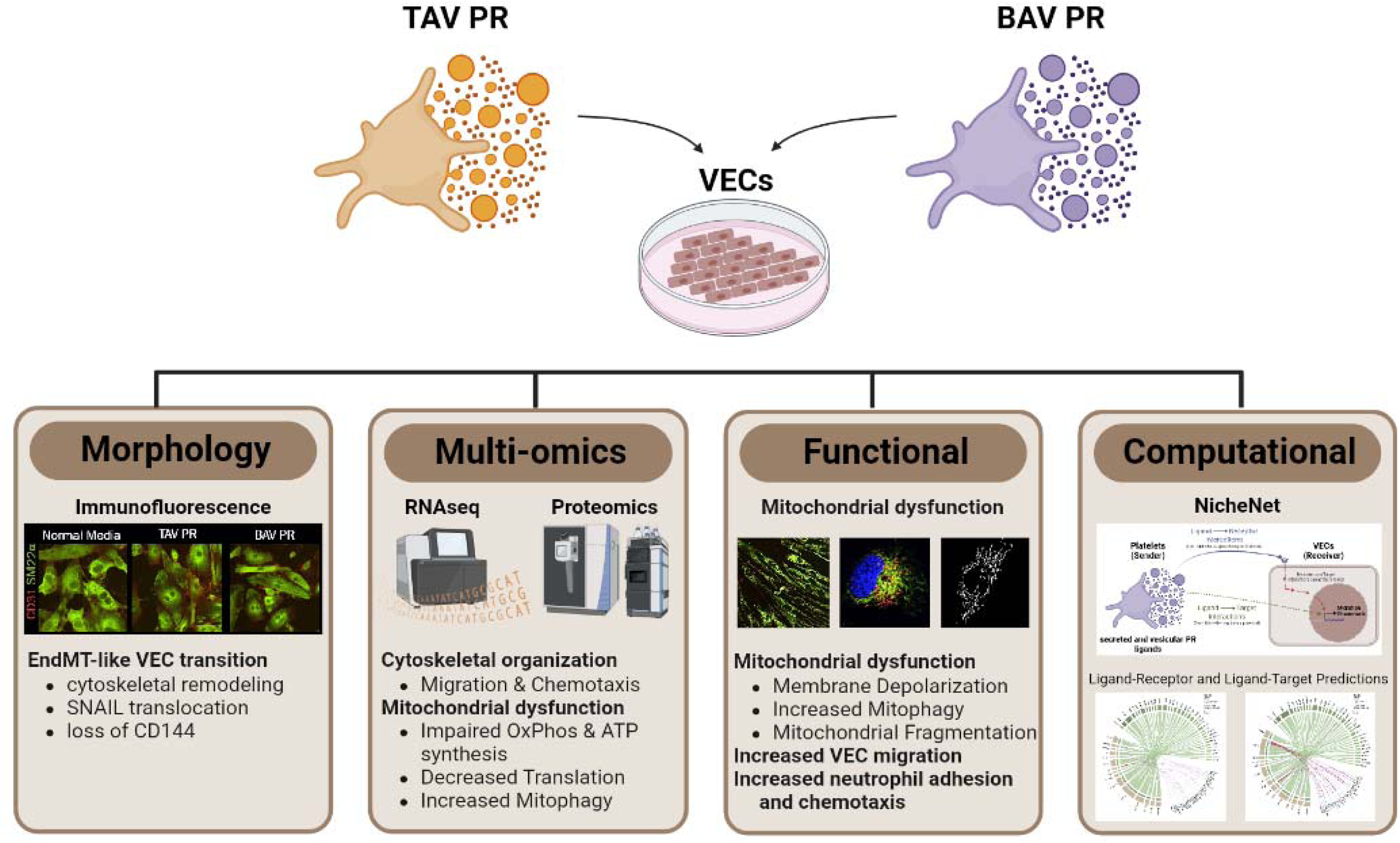

**Translational Perspective:** Bicuspid aortic valve (BAV) confers a substantial risk of premature aortic stenosis, yet no pharmacological therapy to halt disease development exists. Our findings position platelet-endothelial crosstalk as a mechanistic link between the abnormal hemodynamic environment of BAV and valvular endothelial dysfunction. Platelet-derived mediators promoted endothelial remodeling, mitochondrial dysfunction, and a migratory phenotype, with more marked effects in BAV. These observations suggest that platelets actively contribute to the initiation and progression of calcific aortic valve disease rather than solely reflecting advanced stenosis. Targeting shear-dependent platelet-endothelial signaling may therefore provide new opportunities for sought-after disease-modifying therapeutic strategies in (BAV-associated) aortic stenosis.

## 1. Introduction

Calcific aortic valve disease (CAVD) is the most prevalent valvular heart disease and is characterized by progressive inflammation, fibrosis and calcification of the aortic valve leaflets, ultimately leading to aortic stenosis (AS). Despite the global surge in AS prevalence, the paucity of pharmacological therapies to prevent or halt disease progression leave valve replacement as the only therapeutic option.^1^ Rather than a passive consequence of ageing, CAVD is now recognized as an actively regulated process in which pathological hemodynamics drive endothelial dysfunction, inflammation and tissue remodeling.^2^

Valvular endothelial cells (VECs) form the interface between the circulation and valve tissue and are therefore uniquely positioned to translate altered hemodynamic and circulating signals into disease-promoting responses. Under physiological conditions, VECs maintain barrier integrity, nitric oxide signaling and an anti-inflammatory, antithrombotic surface.^3^ During CAVD, disturbed shear stress promotes endothelial activation and dysfunction, including oxidative and inflammatory signaling, leukocyte recruitment, impaired barrier function and endothelial-to-mesenchymal transition (EndMT).^4,5^ These changes disrupt VEC-valvular interstitial cell (VIC) communication and actively promote fibrotic and osteogenic valve remodeling.^6,7^

Emerging evidence suggests that platelets may provide an important link between pathological hemodynamics and endothelial dysfunction in CAVD. Beyond the increased platelet activation induced by abnormal hemodynamic stresses in AS^8,9^, activated platelets accumulate at sites of endothelial disruption in calcified human valves and directly promote osteogenic responses in VICs^10^, highlighting their active role in valvular remodeling. Importantly, platelet activation triggers the release of a complex bioactive secretome, termed the platelet releasate (PR)^11^, containing inflammatory, vasoactive, adhesive and growth-regulatory mediators. These factors can substantially alter endothelial signaling, permeability, migration and leukocyte interactions, providing a mechanism through which hemodynamically activated platelets could influence the valve even without sustained platelet adhesion. Despite extensive evidence for platelet-endothelial crosstalk in vascular disease^12^, the effects of platelet-derived mediators on VECs remain elusive.

This interaction may be particularly important in bicuspid aortic valve (BAV) disease, the most common congenital valve abnormality and strongest non-modifiable risk factor for accelerated AS. BAV geometry generates eccentric flow and abnormal, oscillatory shear patterns that precede severe valve dysfunction and are sufficient to induce VEC activation and maladaptive changes.^13^ Although BAV and tricuspid aortic valve (TAV) VECs exhibit distinct molecular phenotypes and susceptibilities to cellular stress^14,15^, whether platelet-derived signals differentially regulate valvular endothelial function in BAV and TAV remains unidentified. In particular, it remains unknown how the distinct hemodynamic environments associated with valve pathomorphology influence platelet activation and the release of platelet-derived mediators. The direct effects of these functionally distinct platelet releasates on inducing divergent VEC phenotypes remains unclear. Addressing this gap allows to situate circulating platelets at the interface of valve hemodynamics and endothelial dysfunction and provides a potential mechanism through which abnormal flow may be converted into sustained biochemical signaling within the valve. Investigating PR independently of the upstream mechanisms driving platelet activation may therefore provide important insights into the direct contribution of platelet-derived signaling to valvular endothelial dysfunction.

We hypothesized that molecular cargoes differ in PR from patients with BAV and TAV and consequently provoke distinct molecular and functional responses in VECs. To test this hypothesis, we integrated molecular profiling of PR with transcriptomic, proteomic and functional characterization of human VEC responses to BAV- and TAV-derived PR to determine whether platelet-derived signals contribute to morphology-specific endothelial dysfunction, identify mechanisms linking altered hemodynamics, platelet activation and valvular disease progression, and uncover potential therapeutic targets to disrupt this pathogenic crosstalk.

## 2. Methods

A detailed description of all materials and methods is provided in the Supplementary material online.

### 2.1. Patient recruitment, blood sampling, and platelet releasate generation

Patients undergoing surgical aortic valve (AV) replacement as part of their clinical care were prospectively recruited at Brigham and Women’s Hospital (BWH), Boston, MA. The study protocol was approved by the Institutional Review Board and Human Research Committee at BWH (2011P001703 and 2025P001420). All participants provided informed written consent according to the Declaration of Helsinki. Fasting blood samples were collected into acid citrate dextrose A prior to administration of any sedatives on the morning of valve replacement surgery. Platelet releasate (PR) was prepared as previously described^16^ and stored at −80°C until use.

### 2.2. Valve endothelial cell (VEC) isolation and culture

Explanted human aortic valve tissue (n=6 BAV, n=6 TAV) was collected from patients undergoing valve replacement surgery at BWH. VECs were isolated from aortic valve leaflets according to our published protocol.^17^

VECs were cultured in collagen-coated flasks in EGM-2 with 10% FBS and 1% antibiotic (CC-3162) at 37°C in a humidified incubator with 5% CO_2_ and used for experiments between passages 3 and 7. For each experiment, VECs were seeded at 1x10^5^ cells/mL 24 hours prior to treatments. VECs were treated with TGFβ (5 ng/mL) and TNFα (50 ng/mL). For PR treatments, VECs were cultured in EGM-2 supplemented with 2.5% (v/v) PR and 2.5% (v/v) FBS, resulting in a total serum/releasate supplementation of 5%. Control and cytokine-treated VECs were cultured in EGM-2 supplemented with 5% FBS. To prevent medium gelling caused by active coagulation factors in the PR, 0.6 U of heparin were added to all treatment conditions. VECs were treated for 4 days with a media change every 48 hours. Conditioned media was collected on day 4 and stored at -80°C.

### 2.3. RNA sequencing

VECs were lysed with 70 µL Zymo DNA/RNA shield^TM^ (R1 100-50) and submitted to Plasmidsaurus (Eugene, OR, USA) for library preparation and RNA sequencing. After annotation, raw counts were normalized, log2 transformed, and assessed for statistically significant differential expression using edgeR and limma packages. Resulting *p*-values were adjusted for multiple testing using Benjamini–Hochberg (BH) correction. For visualization and subsequent pathway analyses, genes with an absolute log2 fold change >1 and FDR ≤0.05 were considered differentially expressed.

Principal component analysis (PCA) was performed to visualize treatment-associated variation independently of donor-specific effects. Donor effects were removed from the expression matrix, while retaining treatment-associated variation in the model. Genes with zero variance or incomplete expression values were excluded prior to PCA. The proportion of variance explained by each principal component was calculated. Donor correction was performed exclusively for visualization and quality-control analyses; differential expression testing was performed on the uncorrected expression data with donor explicitly incorporated as a covariate in the linear model.

### 2.4. Mass Spectrometry

#### 2.4.1. PR proteomics sample preparation

PR samples were lysed with equal volumes of RIPA buffer for 10 min on ice and the protein concentration was determined by BCA. 10 ug protein were used as input. Mass spectrometry sample preparation was performed using the PreOmics 96 iST-BCT kit (P.O.00099, PreOmics) according to the manufacturer’s instructions.

#### 2.4.2. VEC proteomics sample preparation

VECs were lysed in 50 µL RIPA buffer for 10 min on ice. Protein concentration was determined by BCA. 5 ug protein were used as input. Mass spectrometry sample preparation was performed using the PreOmics 96 iST kit (P.O.00027, PreOmics) according to the manufacturer’s instructions.

#### 2.4.3. Proteomics data acquisition and spectral annotation

Tryptic peptides were analyzed using a quadrupole Orbitrap Exploris 480 coupled to a Vanquish Neo UHPLC system (Thermo Fisher Scientific) as previously described.^18^

The spectra were queried against the UniProt human database (downloaded May 2025; 42,421 entries) using Proteome Discoverer (version 3.3) using the CHIMERYS node with the INFERYS 4.7.0 prediction model (PR) or SEQUEST-HT (VEC). Trypsin/P was set as the digestion enzyme, allowing up to 2 missed cleavages and a minimum peptide length of 7–30 amino acids (1–6 peptide charge). Oxidation (+15.995 Da) of methionine was set as a variable modification, and carbamidomethylation (+57.021 Da) of cysteine was set as a static modification. The fragment mass tolerance was 20ppm. The FDR for PSM confidence was 0.01 and 0.05, for strict and relaxed confidence, respectively. Quantification utilized unique peptides and razor peptides. Razor peptides were used to quantify only the protein with the most identified peptides and not for the other proteins they are contained in. Peptide abundance was normalized by total peptide amount, and each protein intensity was calculated using the sum of its peptide intensities.

The mass spectrometry proteomics data have been deposited to the ProteomeXchange Consortium via the PRIDE partner repository with the dataset identifiers PXD086328 (PR) and PXD086438 (VECs).

### 2.5. Pathway-directed ligand-receptor analysis using NicheNet

To identify BAV PR-derived factors potentially contributing to transcriptional alterations in VECs, pathway-directed ligand activity analysis was performed using NicheNet^19^. Human ligand-receptor interactions and ligand-target regulatory relationships were obtained from the NicheNet v2 framework. Gene sets representing endothelial migration and chemotaxis-associated biological processes were defined using Gene Ontology Biological Process annotations obtained from the Molecular Signatures Database (MSigDB).

Candidate upstream ligands were derived from PR proteomics. Proteins with increased abundance in BAV PR (log2FC>0) that were represented as ligands in the NicheNet ligand-receptor network were retained. To identify ligands potentially contributing to specific endothelial phenotypes, NicheNet ligand activity was calculated against predefined gene sets representing endothelial migration and chemotaxis. Ligands were ranked by their predicted ligand activity. For prioritized ligands, regulatory potential scores from the NicheNet ligand-target matrix were used to identify predicted downstream target genes. Predicted ligand-receptor and ligand-target interactions were subsequently visualized.

### 2.6. Statistical Analysis

All statistical analyses and data visualization were performed in RStudio (version 2024.9.0.375). Statistical analysis for non-omics data sets was performed using a two-sided unpaired Wilcoxon ranksum test followed by Benjamini Hochberg (BH) correction for multiple comparisons. For bulk RNA sequencing, differential gene expression analysis was performed using the edgeR and limma R packages. Genes with low expression were removed using filterByExpr, and library sizes were normalized. Count data were transformed using the voom method to model the mean–variance relationship and generate precision weights for linear modeling. Differential expression was assessed using linear models. Empirical Bayes moderation was applied to fitted models using eBayes. P values were adjusted for multiple testing by BH correction. Genes with an absolute log2 fold change>1 and *p*<0.05 were considered differentially expressed. For proteomics analysis, differential protein abundances were assessed using linear modeling with empirical Bayes moderation of protein-wise variances implemented in the limma package. Statistical significance was determined from the moderated *t*-statistics, with multiple-testing correction applied using a BH false discovery rate (FDR) at 0.05. Genes with an absolute log2 fold change>1 and *p*<0.05 were considered differentially expressed.

## 3. Results

### Platelet releasate composition differs between BAV and TAV

Previous comparative studies investigating shared and divergent disease phenotypes in BAV and TAV predominantly entailed histological, proteomic, and transcriptomic analyses of valve tissue or ascending aortae. We performed the first comparative characterization of platelets and their releasate between BAV and TAV patients. While plasma levels of platelet membrane and α-granule-associated proteins P-selectin (CD62P), platelet factor 4 (PF4), and Thrombospondin-1 (TSP1) were increased in BAV, only TSP1 indicated increased release from BAV following thrombin stimulation (**Figure S1**). Concordantly, thrombin-induced aggregometry demonstrated an attenuated functional response in BAV compared with TAV, characterized by reduced maximal aggregation, aggregation slope, and area under the curve, while the lag time remained unchanged (**Figure S1**). These results point towards a partially preactivated and refractory platelet phenotype, consequently diminishing *ex vivo* responses to thrombin stimulation.

Besides releasing endogenously synthesized proteins, platelets can endocytose circulating mediators into their granules and release them upon activation. Mass spectrometry-based proteomics of PR quantified 1332 proteins (**Table S1**), of which 647 were previously identified in PR from healthy donors (Figure 1A).^20^ As expected, these proteins significantly associated with Reactome pathways implicated in platelet activation and innate immunity (**Figure 1A; Table S2**). Intriguingly, comparison of the previously unidentified PR proteins (n=658) with gene sets from Jensen DISEASES, a database integrating disease–gene associations from biomedical literature, curated resources, and genetic studies, revealed significant enrichment for cardiovascular disease-related terms (**Figure 1B; Table S3**), supporting the potential of platelets as circulating carriers of disease-relevant signatures.

**Figure 1:**
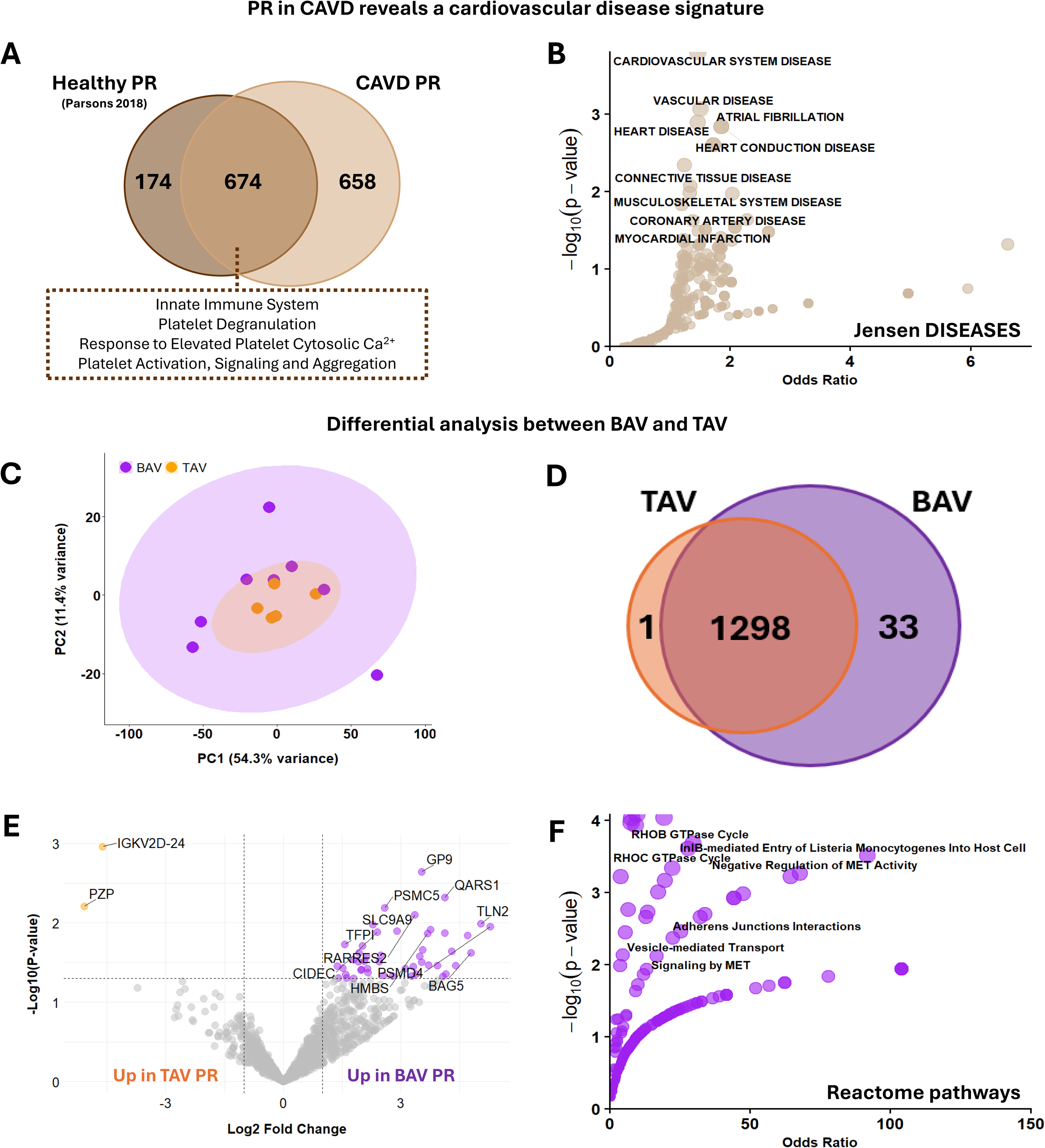
Proteomic profiling reveals valve morphology-specific differences in PR. Mass spectrometry-based proteomics of platelet releasate from CAVD patients with BAV (n=8) and TAV (n=5). (A) Venn diagram of quantified proteins and the overlap with proteins associated with healthy PR. (B) Jensen Diseases enrichment analysis of the previously not reported PR proteins (n=658). (C) Principal Component Analysis (PCA) showing the distribution of BAV and TAV samples, with TAV samples falling within the broader distribution of BAV samples. (D) Venn diagram of the proteins unique to TAV (n=1) or BAV (n=33) or shared between the groups (n=1298). Full list of unique proteins (Supplementary Table 4). (E) Volcano plot of differential enrichment analysis between BAV and TAV PR using a t-test. Proteins highlighted in orange are enriched in TAV (n=2), proteins highlighted in purple and enriched in BAV (n=48, *p*<0.05). (F) Reactome pathway enrichment analysis of proteins uniquely identified in BAV.

Principal component analysis (PCA) revealed substantial overlap between BAV and TAV PR proteomes, with TAV samples forming a distinct cluster within the broader distribution of BAV samples (**Figure 1C**), indicating inter-individual heterogeneity within the BAV cohort while simultaneously reflecting their shared underlying disease etiology. Despite the similarity in the global proteome, 33 and one proteins were uniquely identified in BAV and TAV, respectively (**Figure 1D; Table S4**), with additional 48 and 2 proteins indicating differential enrichment in BAV and TAV PR (**Figure 1E, Table S4**). Reactome pathway analysis on the proteins upregulated and unique in BAV revealed increased enrichment in GO terms associated with Rho GTPase signaling and clathrin-mediated endocytosis (**Table S5**), fundamental regulators of platelet activation and granule release. This is the first report of differences between circulating platelets in BAV and TAV and indicates a hypersecretory platelet phenotype in BAV, providing a potential mechanistic link between altered platelet biology and disease-associated valvular remodeling. Hence, we investigated if these identified molecular differences translate into functionally divergent VEC phenotypes.

### Platelet releasates induce distinct phenotypic changes in VECs

To investigate functional changes, VECs were treated for 4 days with either TAV PR, BAV PR, or TGFβ+TNFα to induce an inflammatory and EndMT-like reference phenotype (**Figure 2A**).^21^ To exclude PR treatment-associated cytotoxicity, VECs were initially exposed to increasing percentages of PR. Cellular metabolic activity was subsequently assessed using an MTT assay and normalized to the normal media control. VECs exposed to up to 2.5% PR maintained MTT metabolic activity (**Figure 2B**); therefore, this concentration was selected for all subsequent experiments.

**Figure 2:**
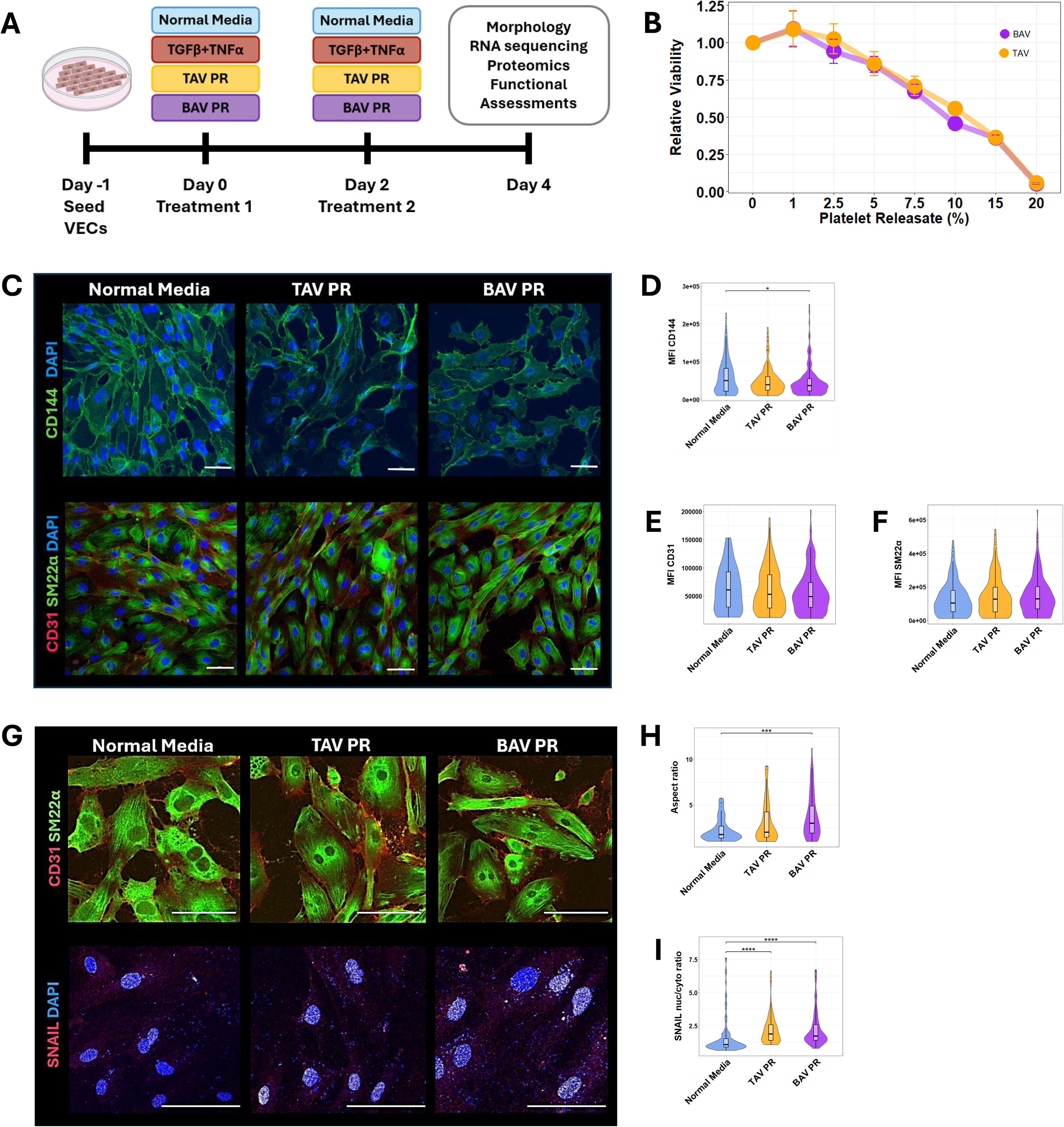
PR induces valve morphology-specific phenotypic changes in VECs. (A) Schematic of the study timelines. (B) MTT assay of n=7 VEC donors (4 BAV, 3 TAV). Data is normalized to normal media control (0% PR) and presented as mean±standard deviation. (C) Representative confocal immunofluorescence images of VECs stained for endothelial markers CD144 (VE-cadherin) and CD31 and mesenchymal marker SM22α at 20x magnification. Nuclei are counterstained with DAPI. Scale bar: 50μm. Quantification of the mean fluorescence intensity (MFI) of (D) CD144, (E) CD31, and (F) SM22α. MFI was normalized to cell count. (G) Representative confocal immunofluorescence images of VECs stained for CD31, SM22α, and SNAIL at 40x magnification. Nuclei are counterstained with DAPI. Scale bar: 50 μm. Quantification of (H) VEC aspect ratio and (I) SNAIL nuclear-to-cytoplasmic ratio. Statistical significance was determined in all instances using a two-sided Wilcoxon ranksum test followed by Benjamini Hochberg correction for multiple comparisons. Data are presented as violin and box-and-whisker plots from n=6 VEC donors (3 BAV, 3 TAV), with the box indicating the interquartile range (IQR), the center line representing the median, and whiskers extending to 1.5xIQR. *\* p<*0.05; ** *p<*0.01; *** *p*<0.001, **** *p*<0.0001.

We first investigated morphological and molecular changes in VECs in response to PR treatment. The expression of endothelial markers CD144 and CD31 and mesenchymal marker SM22α were examined by immunofluorescence (**Figure 2C**). While BAV PR and TGFβ+TNFα induced a loss of CD144 (**Figure 2D; Figure S2**), the expression of CD31 (**Figure 2E**) and SM22α (**Figure 2F**) remained unchanged regardless of the treatment. Moreover, quiescent VECs exhibit a characteristic cobblestone morphology but become elongated as they transition towards a mesenchymal phenotype. This morphological change can be quantified as an increase in aspect ratio. BAV PR and TGFβ+TNFα induced a significant increase in VEC aspect ratio (**Figure 2G and H; Figure S2**), whereas TAV PR had no comparable effect. In addition, translocation of transcription factor SNAIL, an EndMT master regulator, from the cytoplasm into the nucleus validated activation of EndMT processes in VECs. Translocation of SNAIL was increased following PR and TGFβ+TNFα treatment (**Figure 2I; Figure S2**), suggesting that platelet-derived factors promote EndMT-like changes and endothelial dysfunction in VECs, with more pronounced effects elicited by BAV PR. Since canonical EndMT is characterized by a loss of CD31 and increase in SM22α, the observed responses resemble a partial EndMT-like state, prompting further investigation into the molecular mechanisms underlying platelet-induced endothelial dysfunction.

### BAV PR induces distinct transcriptional changes associated with endothelial migration and stress responses

To further delineate the molecular mechanisms underlying these distinct endothelial responses and determine if these reflect canonical EndMT or alternative pathways of endothelial dysfunction, we investigated the transcriptomic profiles of VECs under the different treatment conditions. PCA analysis demonstrated distinct effects of BAV PR on the VEC transcriptome, despite noticeable inter-donor variability (**Figure 3A**). TAV PR-treated samples, on the other hand, clustered more closely with cells maintained in control media. This disparity further emerged when assessing the differentially expressed genes relative to control conditions. BAV PR triggered a distinct set of increased (**Figure 3B; Table S6-7**) and decreased (**Figure 3C; Table S6-7**) genes, whereas TAV PR produced no unique transcriptional changes. Nonetheless, BAV and TAV PR shared eight upregulated and four downregulated genes, including CXCL12 and CXCL6, respectively.

**Figure 3:**
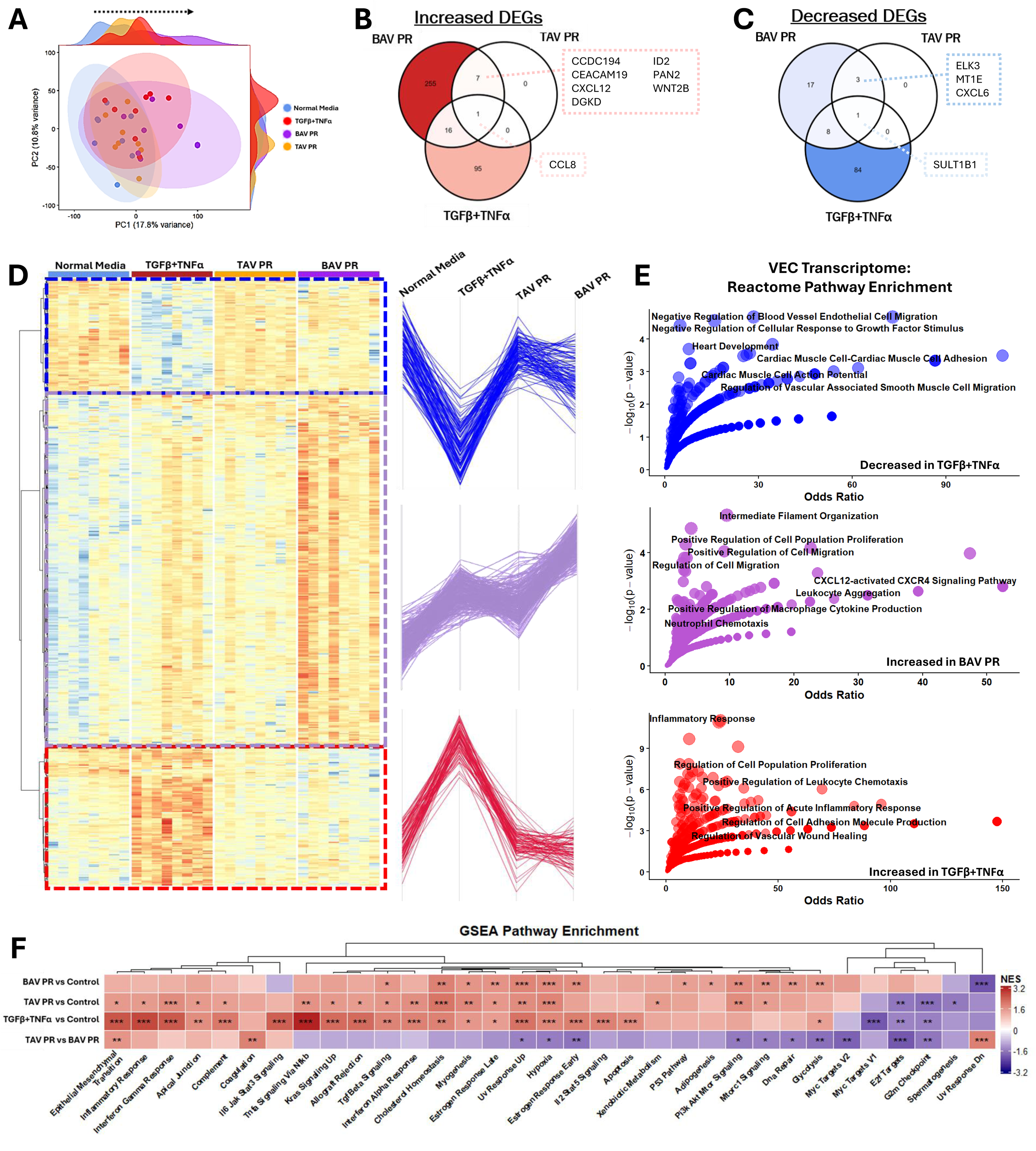
BAV PR promotes a migration-associated transcriptional phenotype in VECs. (A) Principal component analysis (PCA) with density plots of VEC gene expression profiles from n=9 VEC donors (5 BAV, 4 TAV). Venn diagrams of significantly increased (B) or decreased (C) genes between TAV PR-, BAV PR-, and TGFβ+TNFα-treated VEC each compared to normal media control conditions. The full list of genes is presented in Table S6. Pairwise contrasts were evaluated using empirical Bayes-moderated t-tests, with Benjamini–Hochberg correction for multiple testing. (D) Unsupervised hierarchical clustering of the 487 differentially enriched proteins (|log2FC|>1*; q*<0.05) and treatment-dependent gene expression trajectories. (E) Reactome pathway enrichment analysis of genes responsive to TGFβ+TNFα treatment or increased following BAV PR treatment. (F) Unsupervised hierarchical clustering of significantly enriched Hallmark gene sets identified by gene set enrichment analysis (GSEA). Enrichment is presented as the normalized enrichment score (NES), with statistical significance determined from the GSEA null distribution, with Benjamini–Hochberg correction for multiple testing. * *p*<0.05; ** *p*<0.01; *** *p*<0.001; **** *p*<0.0001.

Hierarchical clustering of the 287 differentially expressed genes (cumulative normal media vs treatments) identified distinct gene clusters with treatment-specific expression patterns (**Figure 3D**). As expected, TGFβ+TNFα treatment induced a classical inflammatory endothelial phenotype, characterized by activation of inflammatory pathways alongside suppression of genes involved in the negative regulation of endothelial migration (**Figure 3E; Table S8**). BAV PR-treated VECs, on the other hand, exhibited enrichment of processes associated with cytoskeletal remodeling, including intermediate filament and supramolecular fiber organization, combined with increased regulation of cell migration and proliferation (**Figure 3E; Table S9**), collectively suggesting a transcriptional program indicative of endothelial remodeling and motility.

To assess the relative contributions of treatment conditions and VEC donor variability to gene expression patterns, we performed variance partitioning analysis (**Figure S3**). Consistent with the considerable inter-donor heterogeneity observed by PCA and hierarchical clustering, variance partitioning attributed ∼35% of transcriptomic variability to donor identity and only 5% to the treatment condition. Ranking and clustering the top 100 treatment-associated genes according to the proportion of variance attributed to treatment identified a distinct TGFβ+TNFα-responsive transcriptional signature. This likely reflects the potent and targeted transcriptional responses elicited by these established inflammatory stimuli, in contrast to the more complex and heterogeneous effects of biologically derived platelet mediators. Importantly, this distinction does not diminish the relevance of PR-induced transcriptional changes but rather highlights the fundamentally different nature of these responses, with PR potentially engaging multiple, more subtle signaling pathways that collectively contribute to endothelial dysfunction. To exclude the possibility that VEC origin influenced the response to PR, we compared transcriptional profiles of BAV- and TAV-derived VECs cultured in normal control media (**Figure S4**). PCA revealed complete overlap between VECs of BAV and TAV origin, and differential gene expression analysis identified no significant differences between these two VEC populations, suggesting similar TAV and BAV VEC phenotypes in static culture. Similarly, sample correlation analysis indicated no segregation by valve phenotype, within-group transcriptomic distances were comparable between BAV- and TAV-derived VECs (*p*=0.24), and VEC responses did not differ between VEC origin (**Figure S4**). Although overall transcriptomic variability did not differ significantly between valve phenotypes, potentially meaningful differences in heterogeneity may remain undetected due to the limited sample size.

To further characterize treatment-specific differences in molecular pathway activation, we conducted gene set enrichment analysis (GSEA) (**Figure 3F; Table S10**). Exposure to TGFβ+TNFα induced a prominent inflammatory signature, with positive enrichment of pathways associated with inflammatory and interferon-γ responses, IL6-JAK-STAT3 and TNFα/NFκB signaling, consistent with broad activation of inflammatory and stress-responsive transcriptional programs, while proliferative pathways displayed negative enrichment. In comparison, both BAV and TAV PR were associated with positive enrichment of stress-response pathways implicated in AS, including TGFβ signaling, cholesterol homeostasis, and PI3K-AKT-mTOR-related signaling^22,23^, while inflammatory and immune-associated pathways were solely enriched in TAV PR-treated VECs. In contrast, Epithelial-to-Mesenchymal transition was strongly positively enriched in TAV PR relative to BAV PR. Together, these findings reveal distinct PR-induced transcriptional signatures in VECs, with BAV PR preferentially activating metabolic, growth-related, and stress-response pathways, while TAV PR appears to promote a more inflammatory endothelial phenotype.

### BAV PR-induced proteomic changes associate with mitochondrial dysfunction

We next examined whether these transcriptional changes were also reflected at the proteomic level. Proteomic analysis quantified 3,263 proteins, with high concordance in protein abundance between groups (*r*>0.964) and comparable abundance distributions (**Figure S5**). Consistent with the RNA-sequencing data, PCA revealed divergent proteomic trajectories in VECs exposed to BAV PR and TGFβ+TNFα, whereas protein expression profiles of TAV PR-treated cells overlapped with normal media controls, further validating a distinct VEC response to BAV PR treatment (**Figure 4A**). Consistent with this distinct trajectory, BAV PR provoked more noticeable proteomic changes in VECs than TAV PR when compared with control conditions (**Figure 4B and C; Table S11-12**). Enrichment of GO terms involved in inflammatory processes, endothelial migration, and chemotaxis following BAV PR treatment (**Figure 4B; Table S13**) coincided with coordinated suppression of bioenergetic pathways (**Figure 4C; Table S14**), characterized by impaired mitochondrial energy metabolism and *de novo* metabolic processes. Despite the moderate number of unique protein expression changes, the proteins uniquely decreased in TAV PR-treated VECs represented functionally interconnected components of mitochondrial homeostasis, including mitochondrial ribosomal proteins (MRPL45, MRPL57 and MRPS22), iron–sulfur cluster biogenesis machinery (NFS1 and FXN), and respiratory Complex I components (NDUFS6 and NDUFB8). Further corroborating these findings, Hallmark GSEA revealed a shared PR-associated transcriptional signature characterized by suppression of oxidative phosphorylation, indicating general platelet-mediated disruption of mitochondrial function and a TAV PR-specific induction of interferon responses (**Figure 4D, Table S15**).

**Figure 4:**
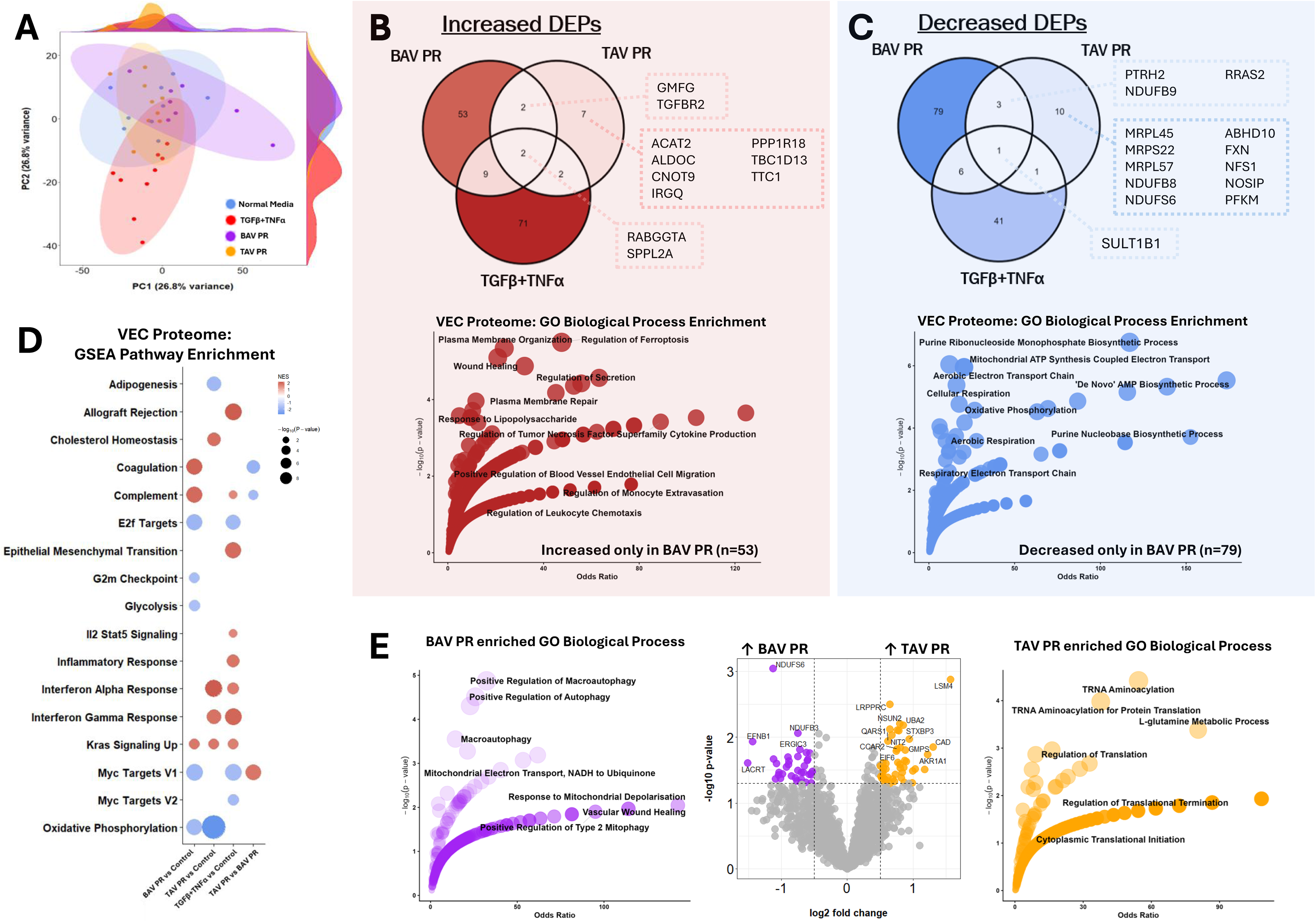
BAV PR induces mitochondrial dysfunction and metabolic remodeling in VECs. (A) Principal component analysis (PCA) with density plots of VEC protein expression profiles from n=9 VEC donors (4 BAV, 5 TAV). (B) Top: Venn diagram of significantly increased proteins between TAV PR, BAV PR-, and TGFβ+TNFα-treated VEC each compared to normal media control conditions. The full list of proteins is presented in Table S11. Pairwise contrasts were evaluated using empirical Bayes-moderated t-tests, with Benjamini–Hochberg correction for multiple testing. Bottom: Gene Ontology (GO) Biological Process enrichment analysis of proteins only increased in BAV PR-treated VECs (n=53). (C) Top: Venn diagram of significantly decreased proteins between TAV PR-, BAV PR-, and TGFβ+TNFα-treated VEC each compared to normal media control conditions. The full list of proteins is presented in Table S11. Pairwise contrasts were evaluated using empirical Bayes-moderated t-tests, with Benjamini-Hochberg correction for multiple testing. Bottom: GO Biological Process enrichment analysis of proteins only decreased in BAV PR-treated VECs (n=79). (D) Bubble plot of the top 5 most significant GSEA Hallmark gene sets per contrast. Bubble color represents NES, while size reflects significance level. Statistical significance is determined from the GSEA null distribution, with Benjamini–Hochberg correction for multiple testing. (E) Direct comparison of BAV PR-vs TAV PR-treated VEC proteomic changes. Volcano plot of differential enrichment analysis using t-test. Highlighted in purple and orange are proteins that meet the significance threshold (|log2FC|>1*; p*<0.05) for differential expression. GO Biological Process enrichment analysis of proteins only increased in BAV PR-treated VECs (n=36, purple) or TAV-treated VECs (n=47, orange).

To further delineate the differential effects of BAV and TAV PR treatment on VECs, we directly compared VEC proteomic changes between the two PR treatments (**Figure 4E; Table S11**). We identified 83 proteins that were differentially enriched (*p*<0.05), with 36 and 47 proteins increased in BAV and TAV, respectively. In line with the identified mitochondrial dysfunction phenotype when compared to control conditions, GO terms differentially enriched in BAV-treated VECs associate with macroautophagy, responses to mitochondrial depolarization, and Type 2 mitophagy (**Figure 4E purple; Table S16**) and may represent an adaptive response, facilitating the clearance of dysfunctional mitochondria under cellular stress. A concomitant increase in GO terms related to translation, tRNA aminoacylation, L-glutamine metabolism and purine biosynthesis in TAV PR-treated VECs relative to BAV (**Figure 4E orange; Table S17**) indicates broad suppression of energy-intensive anabolic processes in BAV, complementing the observed impairment of mitochondrial bioenergetics.

### Molecular signatures induced by PR translate into distinct functional responses in VECs

To determine whether the molecular signatures identified through multi-omics profiling translated into functional changes, we assessed the corresponding cellular phenotypes. Given that our multi-omics profiling converged in pronounced platelet-induced mitochondrial dysfunction and type 2 mitophagy, the process of clearing depolarized mitochondria, we assessed the mitochondrial membrane potential using JC-1. Treatment with BAV PR resulted in significant mitochondrial depolarization, while TAV PR showed a similar but non-significant trend (**Figure 5A, B**). Concomitantly, both PR treatments increased mitophagosome formation, with BAV PR inducing a significantly greater increase than TAV PR (**Figure 5 C, D, Figure S7**), while cellular adenosine triphosphate (ATP) production was similarly impaired by both TAV and BAV PR treatment (**Figure 5E**). Moreover, BAV PR treatment induced mitochondrial fragmentation (**Figure 5 F-H**), collectively indicating platelet-mediated mitochondrial dysfunction and activation of mitophagy-associated responses.

**Figure 5:**
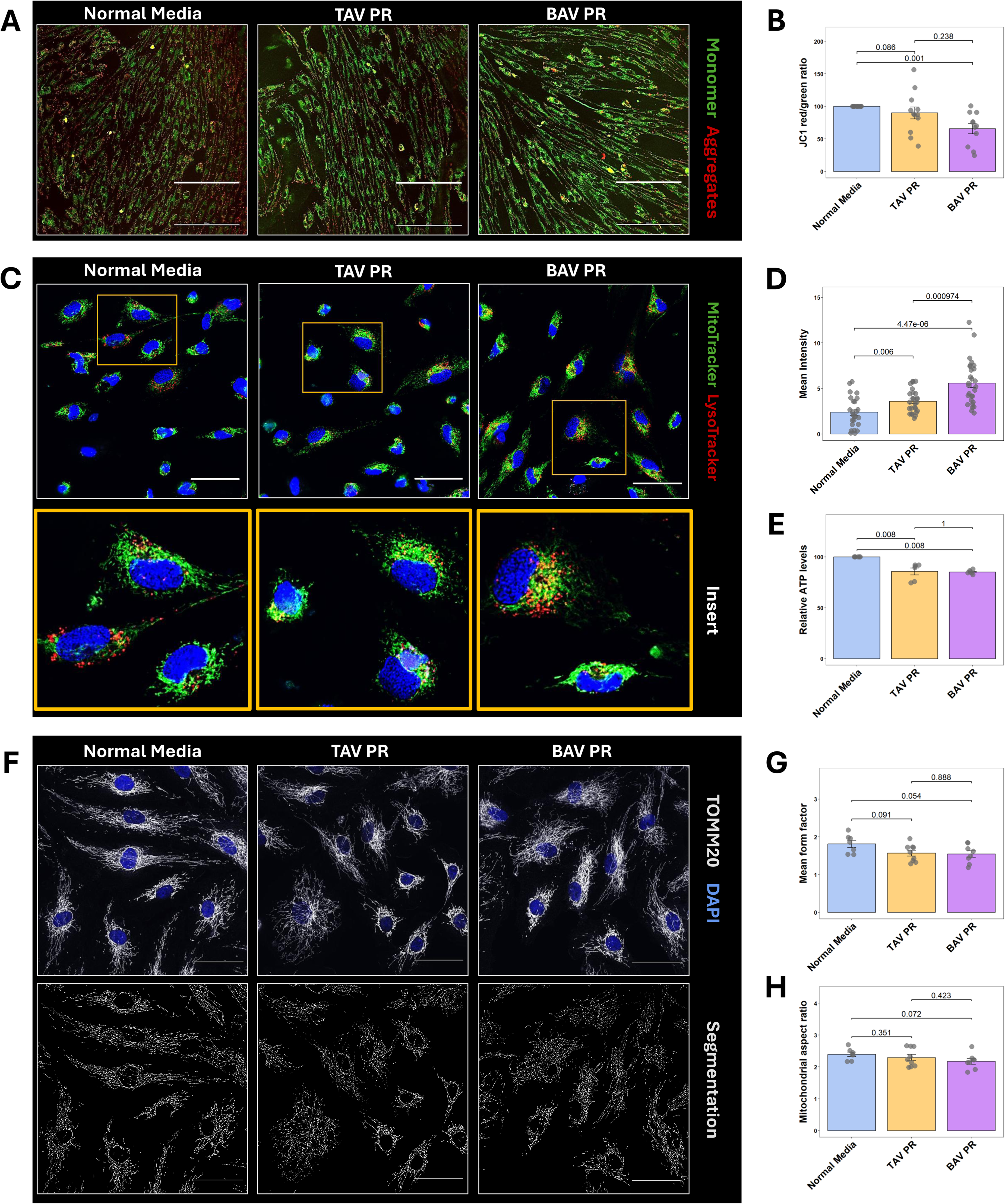
PR treatment induces mitochondrial dysfunction in VECs. (A) Representative confocal images of JC-1 staining of VECs at 20x magnification. (B) Quantification of red-to-green fluorescence ratio. Ratios were normalized to the respective donor matched Normal Media control (n=12). (C) Confocal images of MitoTracker Green and LysoTracker Red staining of VECs at 40x magnification. (D) Mean intensity of MitoTracker Green and LysoTracker Red co-localization area per cell. (E) Quantification of cellular ATP levels normalized to the respective donor-matched Normal Media control (n=6 VEC donors, 3 BAV, 3 TAV). (F) Confocal images of TOMM20 staining and ImageJ-based segmentation of mitochondria. (G) Quantification of the mean mitochondrial form factor per cell. The form factor was calculated as perimeter/(4π*area). (H) Quantification of the mitochondrial aspect ratio. Scale bars: 50 μm.

Beyond metabolic alterations, transcriptomic profiling additionally indicated changes in pathways linked to leukocyte migration, chemotaxis, and wound healing. We therefore investigated whether these molecular changes translated into altered endothelial-immune cell interactions. Given the prominent enrichment of chemotactic pathways and emerging evidence implicating neutrophils in CAVD, we assessed neutrophil adhesion and chemotaxis as functional measures of immune cell recruitment. BAV PR significantly increased adhesion of differentiated HL60 (dHL60) neutrophil-like cells (**Figure 6A, B**) to VECs. HL60 differentiation was confirmed by immunofluorescence staining and flow cytometry (**Figure S8**). Intriguingly, while TAV PR treatment did not significantly increase dHL60 adhesion (*p*=0.078), it elicited a more pronounced chemotactic response (**Figure 6C-E**). This finding is consistent with the increased interferon signaling observed in TAV PR-treated VECs and suggests that TAV PR may promote the release of chemoattractants without substantially enhancing endothelial adhesion molecule expression or stable leukocyte attachment. Lastly, VEC migration in response to endothelial monolayer disruption was significantly enhanced only in BAV PR-treated VECs (**Figure 6F-G**). Collectively, rather than solely representing differences in the magnitude of platelet-mediated endothelial activation, our findings indicate valve morphology-specific molecular and functional responses, with BAV PR preferentially promoting VEC adhesive and migratory phenotypes and TAV PR enhancing neutrophil chemotaxis.

**Figure 6:**
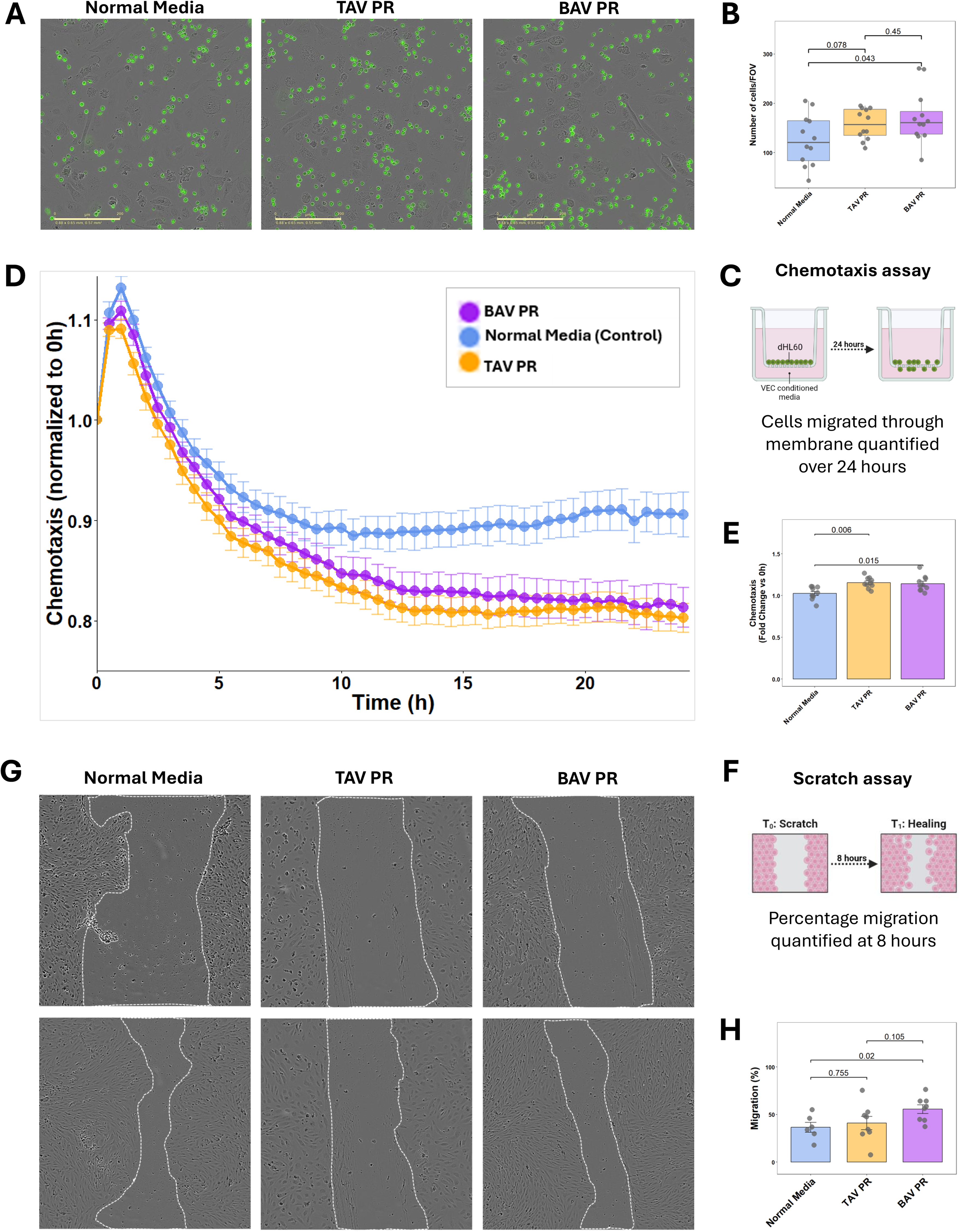
Functional validation of PR-induced VEC phenotypes. (A) Adhesion assay of differentiated HL60 (dHL60) neutrophil-like cells to treated VECs (n=12). dHL60 cells are stained with calcein AM. Scale bar: 200 μm. (B) Quantification of number of adhered dHL60 cells (quantified as number of green objects per field of view (FOV)). (C) Schematic of chemotaxis assay. (D) Chemotaxis assay time course and (E) quantification of dHL60 neutrophil-like cells towards VEC conditioned media (n=12). The cell surface area in the top chamber was normalized by the initial cell area and plotted in a 24-hour time window. The fold change was calculated by the area under the curve for each group. (F) Schematic of the VEC migration assay. (G) Representative brightfield images of wound area at 10x magnification and (H) quantification of migration at 8h normalized to corresponding sample at 0h (n=6-8 VEC donors). Statistical significance was determined in all instances using a two-sided Wilcoxon ranksum test followed by Benjamini Hochberg correction for multiple comparisons. Data are presented as mean±standard deviation or box-and-whisker plots, with the box indicating the interquartile range (IQR), the center line representing the median, and whiskers extending to 1.5xIQR. Individual points represent individual donors.

### PR-derived ligands are predicted to regulate VEC migration and chemotaxis through overlapping and response-specific signaling networks

To identify potential platelet-derived mediators specifically underlying the BAV PR-mediated effects on VEC migration and chemotaxis, we integrated PR proteomics with VEC transcriptomics using NicheNet (**Figure 7A**).

**Figure 7:**
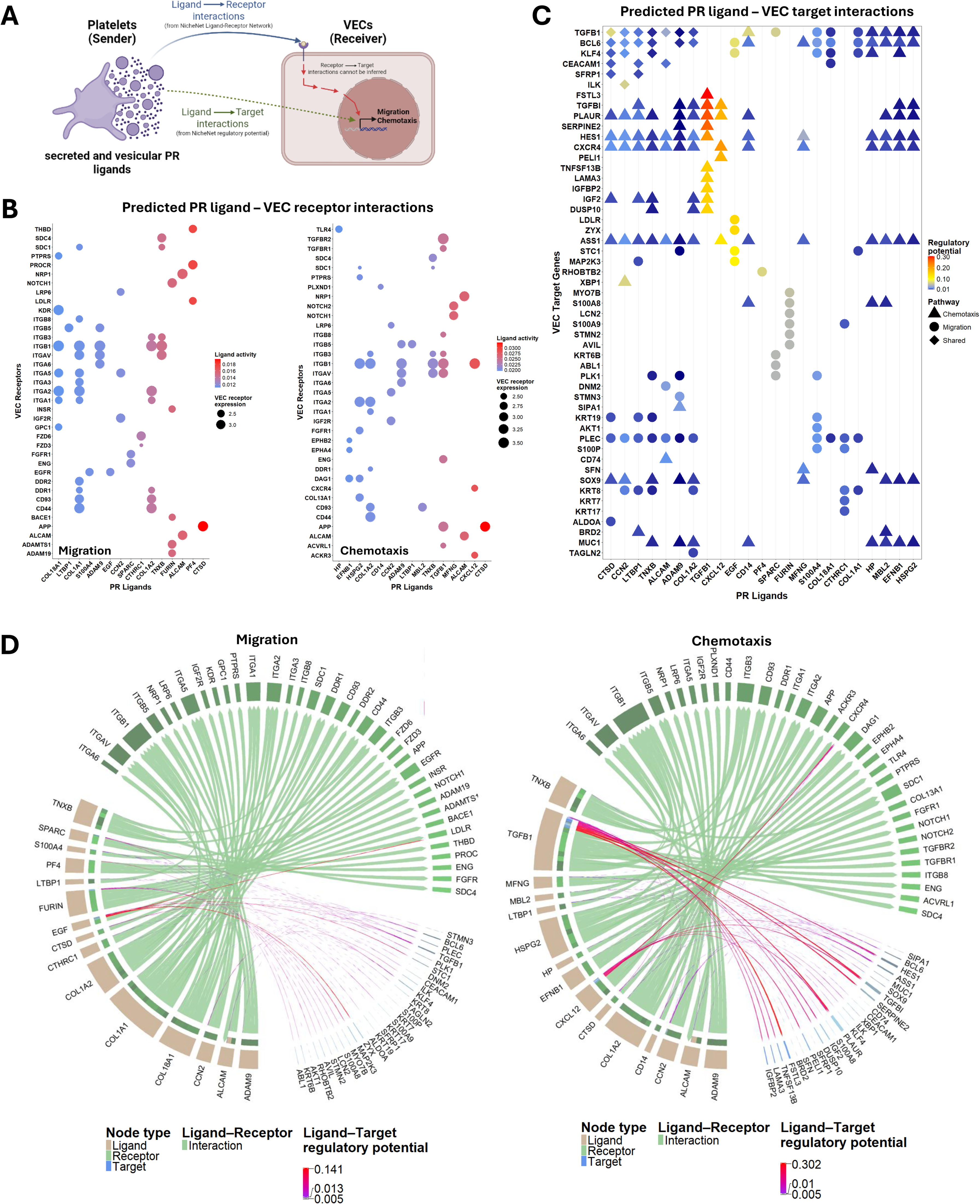
NicheNet analysis identifies PR-derived signaling networks associated with endothelial migration and chemotaxis. (A) Schematic representing the predicted interactions between PR ligands and VEC receptors and targets. (B) Predicted ligand–receptor interactions for migration and chemotaxis, restricted to receptors expressed in VECs. (C) Predicted ligand-target interactions across migration- and chemotaxis-associated gene sets. Ligand-target interaction weights represent NicheNet regulatory potential and do not imply direct or receptor-specific regulation. (D) Chord diagrams depicting the predicted signaling networks associated with migration and chemotaxis, integrating PR-derived ligands with expressed VEC receptors and predicted downstream target genes. Ligand-receptor connections represent interactions supported by the NicheNet ligand-receptor network, whereas ligand-target connections represent predicted regulatory relationships based on NicheNet ligand-target prior knowledge.

Ligand-receptor analysis identified distinct but overlapping BAV PR-VEC communication networks for VEC migration and chemotaxis responses (**Figure 7B, Table S18**). Extracellular matrix (ECM)-integrin signaling emerged as a recurring pattern, with collagen-related ligands, including COL1A1, COL1A2 and COL18A1 and TNXB (tenascin-X), predicted to interact with multiple VEC integrin receptors, suggesting a shared contribution of ECM-sensing and adhesion pathways to BAV PR-induced endothelial responses. Additionally, CTSD (cathepsin D) emerged as a prominent ligand across both migration and chemotaxis networks, with Amyloid Precursor Protein (APP) identified as a predicted endothelial receptor. The CTSD-APP axis has been linked to endothelial homeostasis and amyloid processing^24^, amyloid-like deposits have been identified in calcified aortic valves^25,26^, and APP is an EV-associated driver of VIC calcification^27^, providing disease-specific relevance to this predicted interaction.

Migration-associated signaling additionally included interactions of PF4 with thrombomodulin (THBD) and Protein C receptor (PROCR) (**Figure 7B**). The PF4-THBD-PROCR axis may represent a context-dependent regulatory mechanism balancing endothelial protection and thromboinflammation. Physiological PF4 release supports local activated protein C (APC) generation under homeostatic conditions, thereby reinforcing anticoagulant and endothelial-protective signaling. In contrast, excessive PF4 release during severe vascular stress may promote APC resistance, shifting this balance toward inflammatory endothelial activation, loss of barrier integrity, and a procoagulant state^28^, that further promoted cytoskeletal remodeling through Rho GTPases (**Figure 7C**). The chemotaxis network, on the other hand, identified several ligands implicated in directional migration and endothelial signaling (**Figure 7B**), including TGFB1, CXCL12, and manic fringe (MFNG), with predicted signaling through TGFBR1/2, CXCR4, and NOTCH1/2, respectively, pathways with well-established roles in BAV-associated AS.^29–31^

Ligand-target analysis validated coordinated effects of multiple BAV PR ligands on key endothelial pathways (**Figure 7C, Table S19**). CXCR4, Hairy and Enhancer of Split-1 (HES1), and urokinase plasminogen activator receptor (PLAUR) showed particularly comprehensive ligand connectivity, implicating chemotactic signaling, ECM degradation, and cell-matrix interactions as prominent components of the PR response. Several PR ligands were also linked to TGFB1 and Krüppel-like factor 4 (KLF4), integrating signals involved in shear sensing, inflammatory responses and vascular remodeling. Previous studies positioned KLF4 upstream of TNXB, demonstrating that endothelial KLF4 induces TNXB expression, which in turn restrains TGFβ signaling and EndMT in atherosclerosis through direct interaction with TGFβ^32^. Moreover, TNXB is markedly depleted in calcified aortic valves^33^, potentially pointing towards an interconnected TNXB-KLF4 regulatory axis, in which platelet-derived TNXB may influence KLF4-associated endothelial responses, while KLF4-dependent TNXB expression could provide a feedback mechanism restraining TGFβ signaling and VEC phenotypic remodeling. Additional highly connected targets included plectin (PLEC), a recently identified genetic risk locus for AS^34^, as well as SOX9 and Argininosuccinic synthase 1 (ASS1), drivers of valvular and vascular calcification.^35,36^ Integration of ligand-receptor and ligand-target relationships into a combined network reinforced these recurring patterns (**Figure 7D**). Collectively, these findings identify candidate PR-derived signals capable of contributing to the enhanced migratory and chemotactic VEC phenotype and suggest that these responses arise from a shared core signaling program accompanied by response-specific regulatory networks.

## 4. Discussion

Although BAV- and TAV-associated AS exhibit distinct pathophysiological features, previous comparative studies have largely focused on differences in valvular tissue morphology and resident cellular phenotypes^14,37^, overlooking the potential contribution of circulating factors to disease progression. Platelets, traditionally recognized for their role in hemostasis and thrombosis, are increasingly implicated as active mediators of inflammation, intercellular communication, and tissue remodeling. However, their contribution to valve morphology-specific endothelial dysfunction remains poorly understood. In this study, we provide the first comprehensive characterization of the molecular and functional effects of platelet-derived mediators on VECs, revealing distinct platelet–endothelial signaling responses associated with BAV and TAV. Our findings establish platelet-derived signals as potential contributors to valvular disease pathogenesis and identify molecular pathways that may offer new opportunities for targeted therapeutic intervention. By integrating multi-omics profiling with molecular and functional characterization of VECs and computational signaling analysis, we demonstrated that the PR composition and its downstream endothelial effects differ between patients with BAVs and TAVs, with BAV PR eliciting more pronounced molecular and functional alterations. Importantly, the delineated transcriptional and proteomic signatures reflected altered endothelial migration and leukocyte recruitment, linking platelet-derived molecular signals to functional endothelial phenotypes. Computational signaling analysis further identified candidate platelet-VEC signaling networks that may mediate these responses synergistically and represent personalized therapeutic targets to mitigate endothelial dysfunction and slow disease progression, particularly in BAV.

BAV PR contained a unique repertoire of proteins associated with increased vesicle transport and release, indicating a preactivated but hypersecretory platelet phenotype, despite attenuated aggregation responses. In line with these findings, plasma levels of platelet activation markers increased, and we recently reported increased levels of, among others, platelet-derived extracellular vesicles (EVs) in BAV plasma^38^, substantiating *in vivo* platelet activation in this patient cohort. Intriguingly, mean platelet volume (MPV), a surrogate marker for platelet reactivity, was increased in young, healthy individuals with BAV compared with TAV^39^, suggesting that alterations in platelet phenotype may precede clinically overt valve disease. In contrast, MPV did not differ in our disease cohort (**Table S20**), potentially reflecting the advanced disease stage and comparable hemodynamic burden. In support of the concept that platelets dynamically adapt to changes in their hemodynamic environment across a disease spectrum, MPV decreased 7 days post valve replacement.^40^ Consequently, platelet phenotypes may not represent a static feature of BAV or AS, but rather dynamically reflect the prevailing hemodynamic environment and its trajectory during disease progression.

Beyond differences in PR composition, the biological relevance of platelet-VEC crosstalk lies in its capacity to alter endothelial phenotype. Both BAV and TAV PR induced shared transcriptional changes, including upregulation of CXCL12 and CCL8 and downregulation of CXCL6, suggesting common platelet-mediated modulation of endothelial inflammatory signaling. Although these chemokines have been implicated in AS pathophysiology,^41–44^ their differential regulation highlights the context-dependent effects of platelet-derived mediators. Supporting a role for platelet-mediated inflammation in AS, increased plasma levels of platelet-derived MIF have been associated with accelerated disease progression and platelet infiltration of calcified valves correlates with immune-cell accumulation.^45^ Moreover, platelet-derived CXCL12 can promote endothelial activation and immune-cell recruitment, potentially establishing feed-forward inflammatory signaling.^46,47^ Together, these findings suggest that platelet-derived mediators actively shape the valvular inflammatory microenvironment, promoting endothelial responses that may contribute to AS progression irrespective of valve morphology.

Beyond these shared responses, the distinct composition of BAV PR translated into a more distinct molecular and functional VEC phenotype. EndMT represents a well-established early hallmark of AS^5^ and preliminary evidence situates activated platelets and platelet EVs as inducers of EndMT.^48,49^ Consistently, BAV PR induced an increase in VEC aspect ratio that reflects a shift from the characteristic cobblestone endothelial morphology towards an elongated cellular phenotype. Together with reduced CD144 expression and increased nuclear translocation of SNAIL, these changes suggest an early or incomplete EndMT, rather than a fully established mesenchymal phenotype. Given that EndMT is a dynamic process involving a spectrum of intermediate cellular states^5^, our findings may reflect an initial stage of this transition that could progress with prolonged exposure to platelet-derived mediators.

While representing a general platelet-mediated response, BAV PR induced more definite metabolic perturbations, with converging evidence pointing towards mitochondrial dysfunction. Multi-omics profiling of stenotic human BAV tissue has previously implicated mitochondrial dysfunction, altered oxidative phosphorylation and oxidative stress in BAV-associated AS^50^, while primary VECs isolated from BAV leaflets exhibit increased susceptibility to oxidative stress.^15^ Our findings extend these observations by identifying platelet-derived mediators as a potential extracellular trigger of mitochondrial dysfunction in VECs, irrespective of the valve morphology origin of the VECs. The combination of reduced cellular ATP, mitochondrial depolarization and fragmentation, and increased mitophagosome formation despite enhanced migration following BAV PR exposure further suggests substantial metabolic reprogramming of VECs. Consistent with our observations, platelet lysate depleted intracellular ATP levels^51^ and PR from patients with myocardial infarction induced mitochondrial dysfunction^52^ in vascular ECs. In line with a potential compensatory response to impaired mitochondrial energy production, we identified enrichment of glycolysis at the transcriptomic level, with a concurrent reduction in glycolytic pathways at the proteomic level, indicating that transcriptional activation may not translate into increased glycolytic capacity. Coherently, BAV PR also reduced pathways associated with *de novo* protein synthesis and translation, which could limit the translation of glycolytic transcripts into functional proteins and contribute to this RNA-protein discordance. While these processes may initially represent an adaptive response to mitochondrial dysfunction, persistent mitochondrial fragmentation and dysregulated mitochondrial turnover could ultimately compromise mitochondrial integrity and cellular energy homeostasis. Together, these findings suggest that platelet-derived mediators, particularly those associated with BAV, may contribute to endothelial dysfunction through disturbances in mitochondrial dynamics, quality control, and bioenergetic regulation.

Lastly, a potential importance of hemodynamic forces in modulating platelet biology is supported by clinical observations in severe AS, where abnormalities in high-shear platelet function and loss of high-molecular-weight von Willebrand factor (VWF) multimers correlate with stenosis severity and are rapidly reversed following valve replacement.^9,53^ Although these alterations predominantly reflect shear-dependent VWF proteolysis rather than platelet activation, their immediate normalization following correction of the hemodynamic disturbances underscores the dynamic influence of pathological shear on platelet-VWF biology. Given the severely altered hemodynamic environment imposed by BAV morphology that precedes clinically significant stenosis, our findings raise the possibility that platelet activation in BAV may not solely reflect conventional agonist-mediated signaling but may additionally arise from direct platelet mechanotransduction. Pathological shear stress can activate platelets through mechanosensitive pathways involving VWF-CD42b (also known as GPIbα), integrin αIIbβ3^54^ and Piezo1^55^, pathways which remain active despite conventional antiplatelet therapy.^56–60^ Selective disruption of shear-dependent VWF-GPIbα platelet interactions may therefore represent an unexplored strategy to attenuate platelet-mediated endothelial dysfunction in BAV while avoiding broad suppression of physiological platelet activation. However, whether selective inhibition of platelet mechanotransduction can modify the initiation or progression of BAV-associated AS needs to be established.

Several limitations should be considered when interpreting our findings. First, the *in vitro* PR-VEC model cannot fully recapitulate the complex valvular microenvironment, where endothelial cells are simultaneously exposed to hemodynamic forces and interactions with VICs, immune cells, ECM, and circulating factors. Moreover, PR represents a complex mixture of mediators, and while computational ligand-receptor and ligand-target analyses identified candidate signaling mechanisms, these predictions require experimental validation to establish causal mediator-receptor-target relationships. Second, while mitochondrial membrane depolarization, fragmentation, mitophagosome formation, reduced cellular ATP levels, and proteomic alterations collectively support impaired mitochondrial homeostasis following BAV PR exposure, mitochondrial respiration and glycolytic flux were not directly assessed. Consequently, the contribution of mitochondrial dysfunction and potential compensatory metabolic reprogramming to the observed phenotype remains to be established. Finally, while our findings support distinct platelet-mediated effects in BAV and are consistent with a potential contribution of the abnormal BAV hemodynamic environment, platelet mechanotransduction was not directly investigated. As both blood and valve samples were obtained from patients with established AS, the present study cannot determine whether the BAV-specific platelet phenotype precedes and contributes to disease progression or develops secondary to the altered hemodynamics of established valve disease.

In conclusion, our findings identify platelet-VEC crosstalk as a novel component of CAVD pathobiology and reveal important differences associated with valve morphology. BAV and TAV were characterized by distinct PR profiles that translated into differential molecular and functional responses in human VECs. Our results support a novel model in which valve morphology-specific hemodynamic environments shape platelet activity and PR composition, thereby providing a circulating mechanism through which abnormal flow may propagate endothelial dysfunction and contribute to the accelerated CAVD phenotype associated with BAV. Targeting platelet-endothelial signaling, particularly upstream shear-dependent platelet activation or specific PR-mediated signaling pathways, may therefore provide novel avenues to disrupt early disease-instigating mechanisms before irreversible valvular remodeling and calcification occur.

## 5. Acknowledgements

L.W. is supported by the European Union’s Horizon Europe Research and Innovation Program under the Marie Skłodowska-Curie Actions (101203749). M.A. and E.A. are supported by the National Institutes of Health R01HL174066 and research grant from Kowa Company, Japan. E.A. is supported by the Leducq Foundation PRIMA network (22RAF02), a research grant from Pfizer (GR 1000131), and the American Heart Association Strategically Focused Research Network (26SFRNCVHD1592182).

