## Supplementary Figures for "Platelet-endothelial crosstalk drives distinct endothelial dysfunction in bicuspid aortic valve disease"

### Slide 1
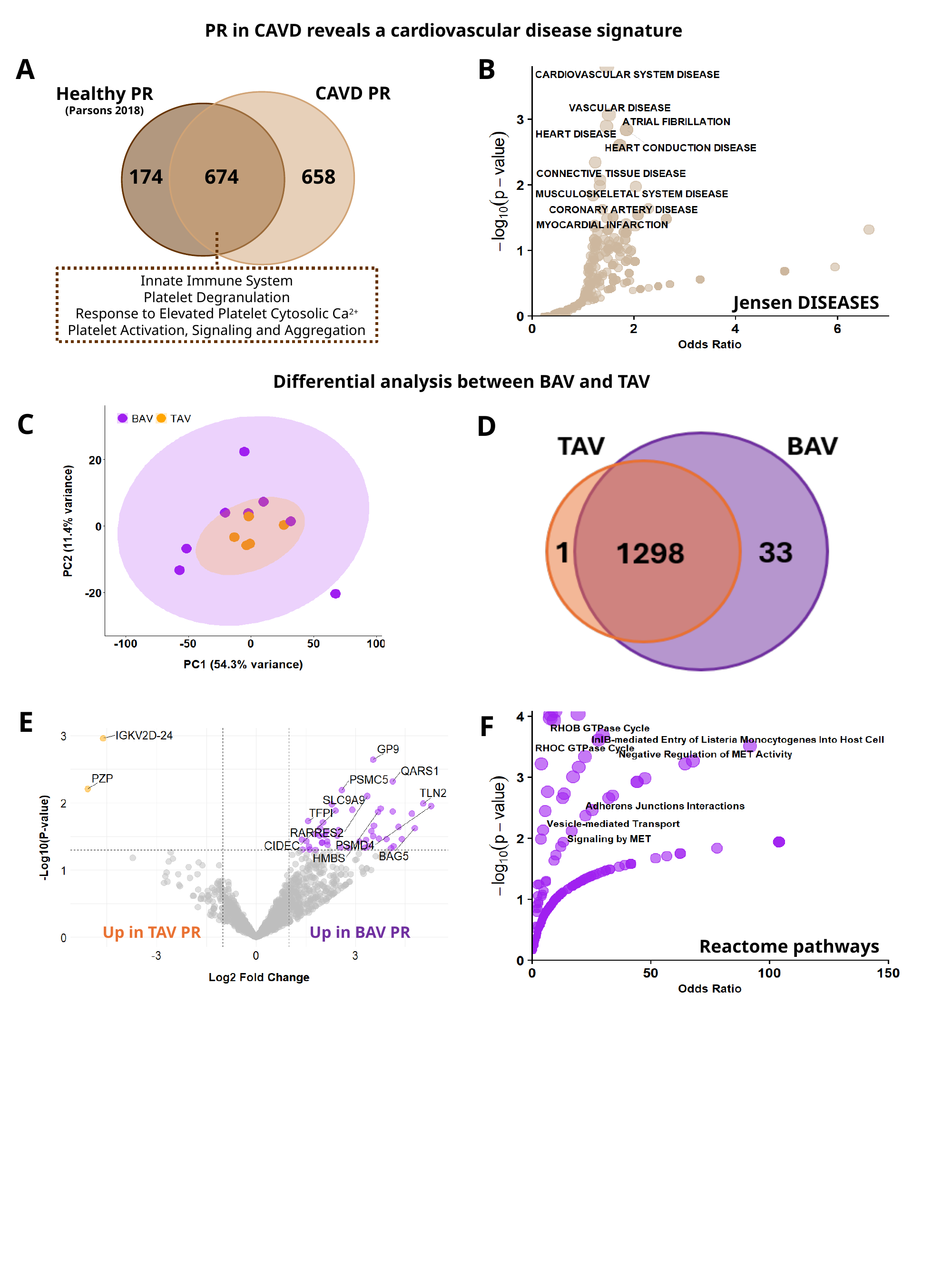

PR in CAVD reveals a cardiovascular disease signature
A
B
CAVD PR
Healthy PR
(Parsons 2018)
674
658
174
Innate Immune System
Platelet Degranulation
Response to Elevated Platelet Cytosolic Ca^2+^
Platelet Activation, Signaling and Aggregation
Jensen DISEASES
Differential analysis between BAV and TAV
C
D
E
F
Up in TAV PR
Up in BAV PR
Reactome pathways

### Slide 2
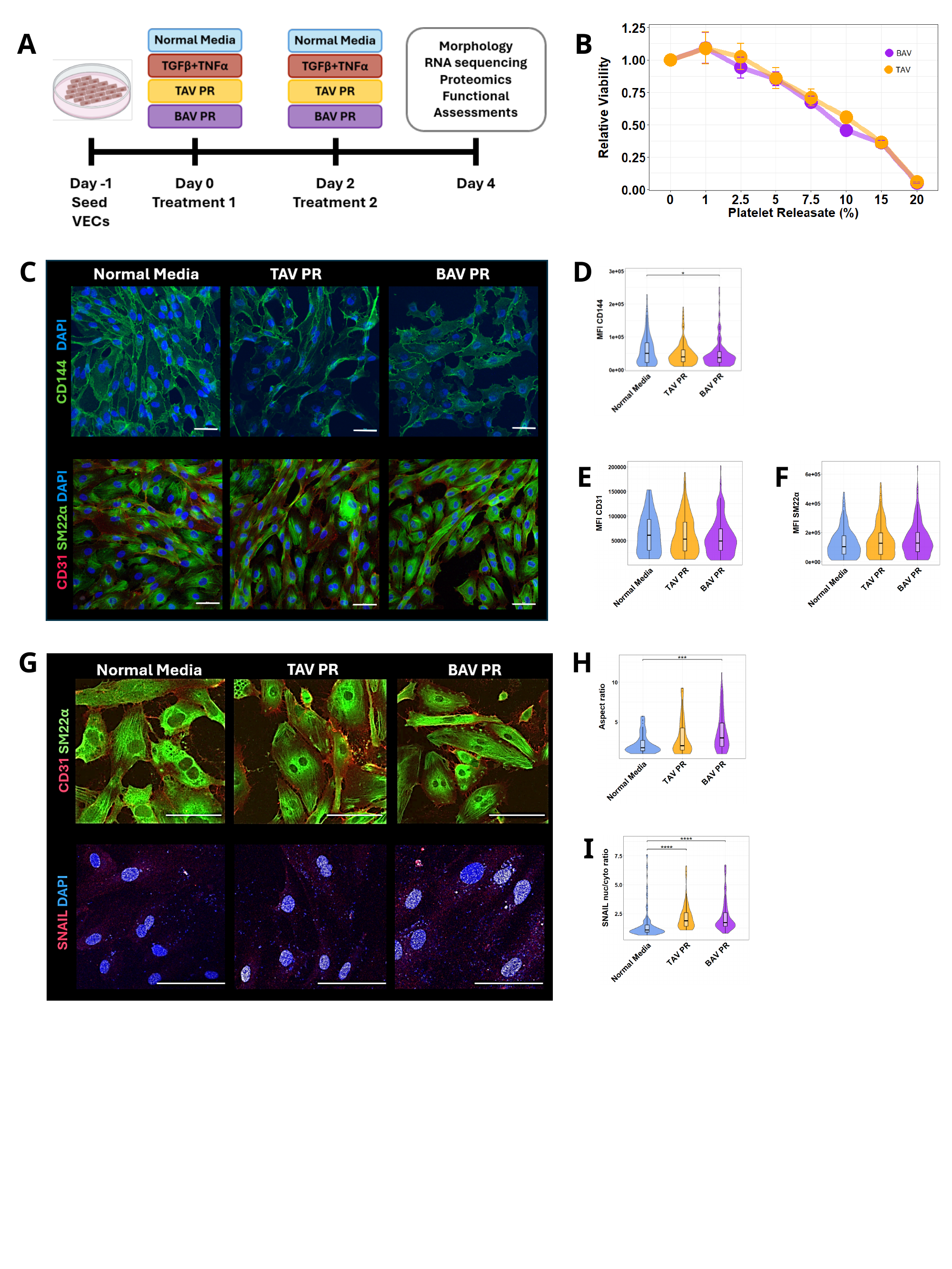

B
A
C
D
E
F
G
H
I

### Slide 3
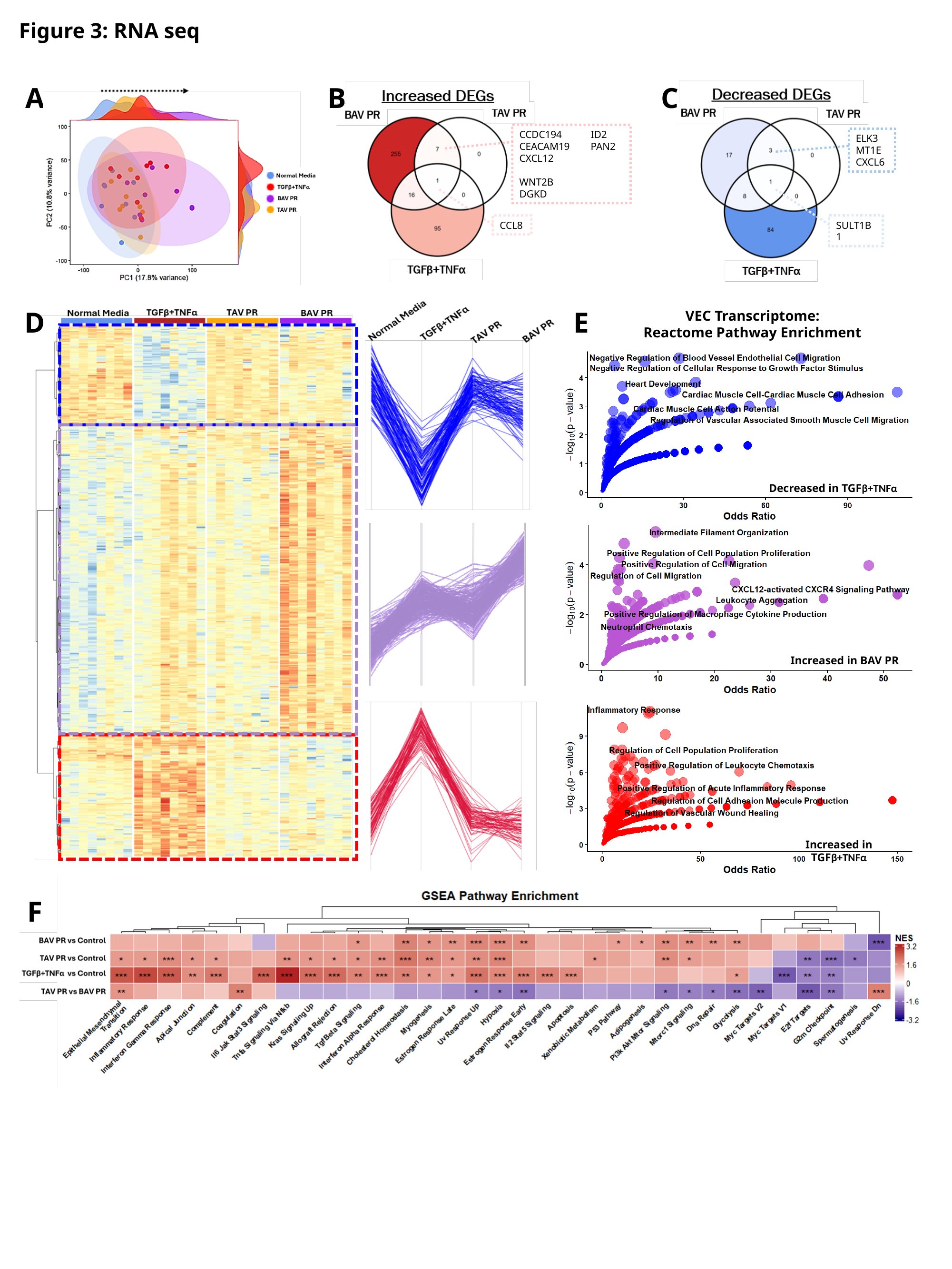

Figure 3: RNA seq
A
B
C
CCDC194	ID2
CEACAM19	PAN2
CXCL12		WNT2B
DGKD
ELK3
MT1E
CXCL6
CCL8
SULT1B1
D
E
VEC Transcriptome:
Reactome Pathway Enrichment
Decreased in TGFβ+TNFα
Increased in BAV PR
Increased in TGFβ+TNFα
F

### Slide 4
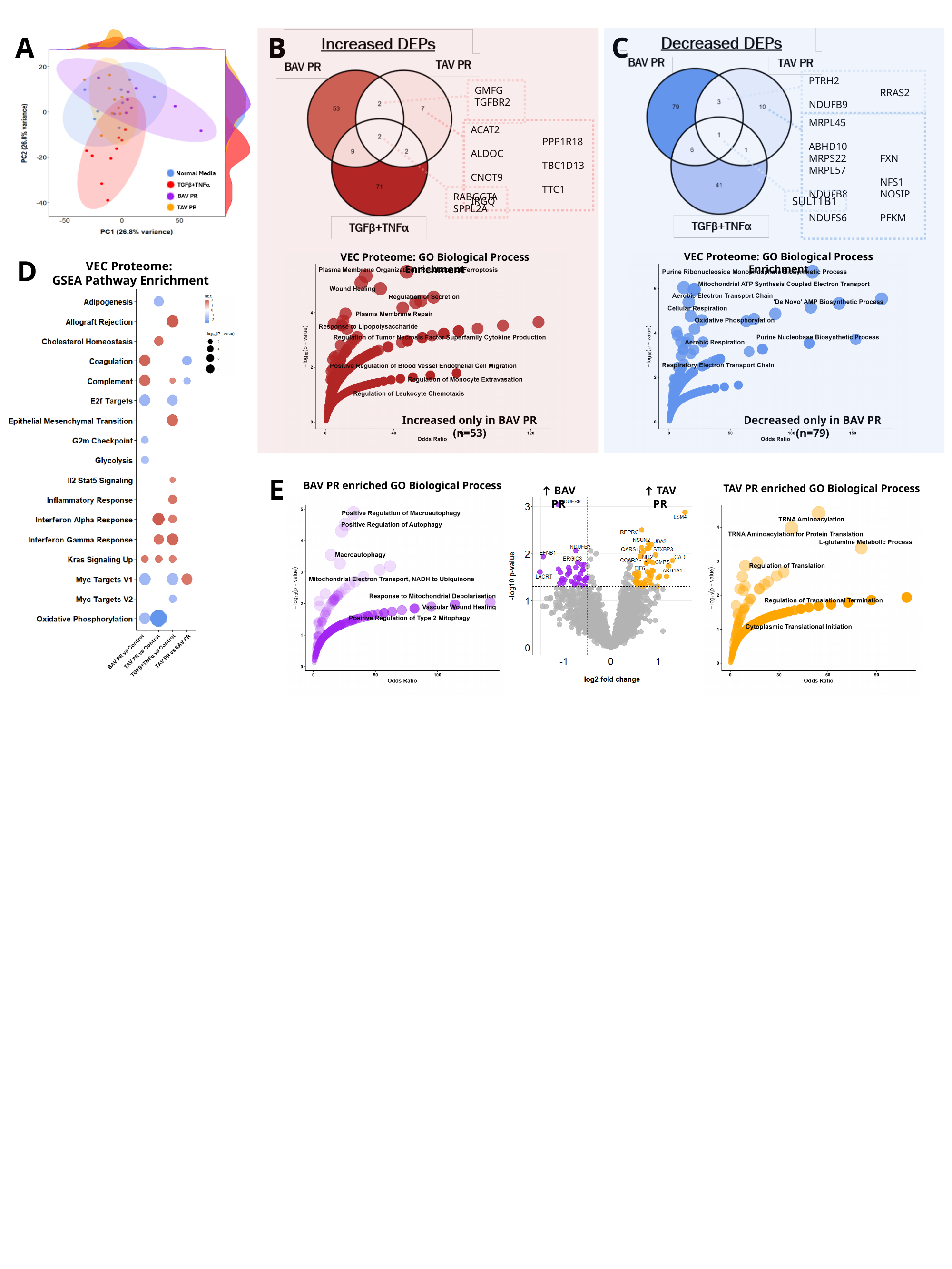

A
B
C
PTRH2		RRAS2
NDUFB9
MRPL45		ABHD10
MRPS22	FXN
MRPL57		NFS1
NDUFB8	NOSIP
NDUFS6	PFKM
SULT1B1
B
GMFG
TGFBR2
ACAT2		PPP1R18
ALDOC		TBC1D13
CNOT9		TTC1
IRGQ
RABGGTA
SPPL2A
VEC Proteome: GO Biological Process Enrichment
VEC Proteome: GO Biological Process Enrichment
D
VEC Proteome:
GSEA Pathway Enrichment
Increased only in BAV PR (n=53)
Decreased only in BAV PR (n=79)
E
BAV PR enriched GO Biological Process
TAV PR enriched GO Biological Process
↑ BAV PR
↑ TAV PR

### Slide 5
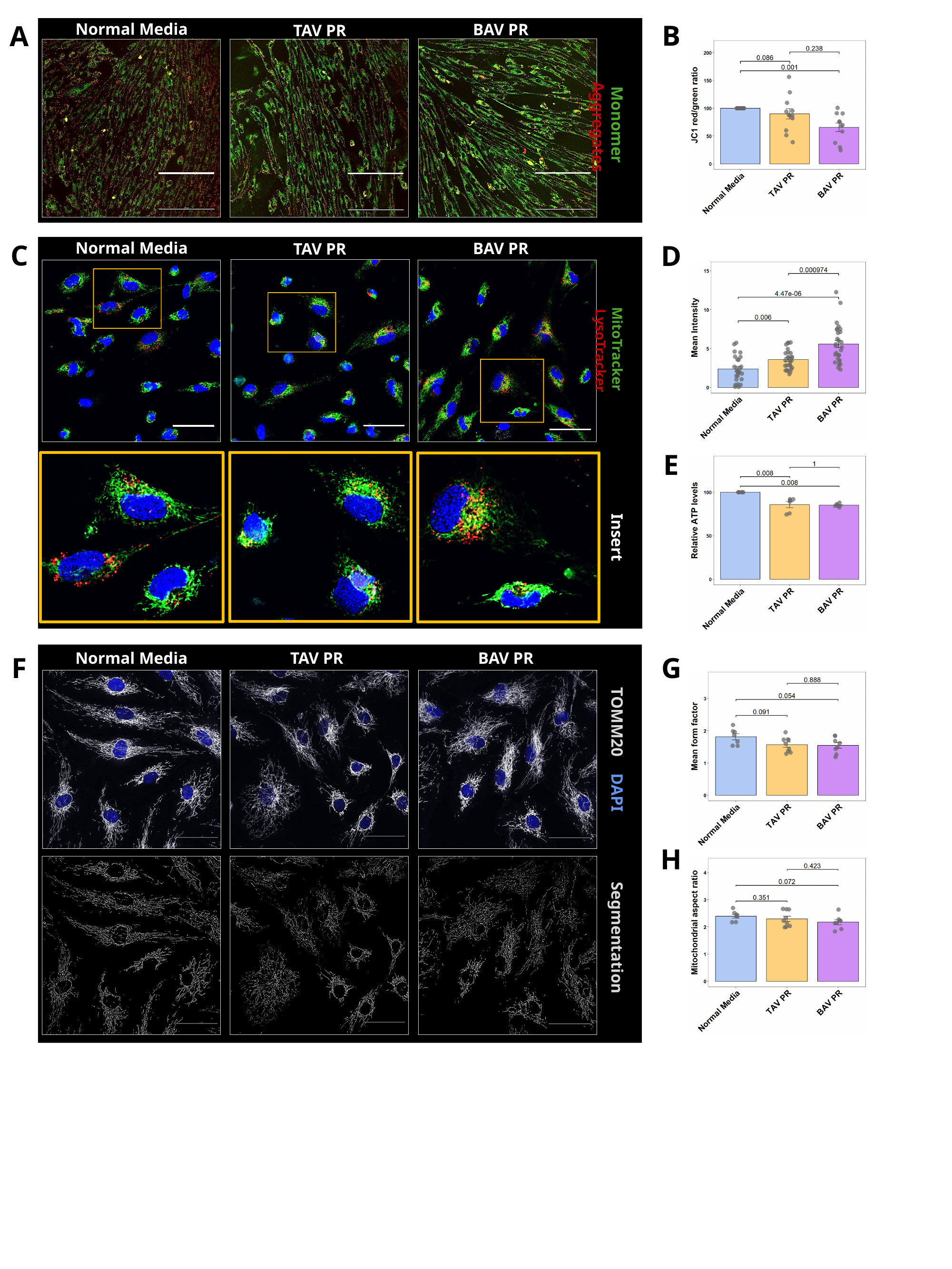

B
A
Normal Media
BAV PR
TAV PR
Monomer Aggregates
Normal Media
BAV PR
TAV PR
C
D
MitoTracker LysoTracker
E
Insert
Normal Media
TAV PR
BAV PR
TOMM20 DAPI
Segmentation
F
G
H

### Slide 6
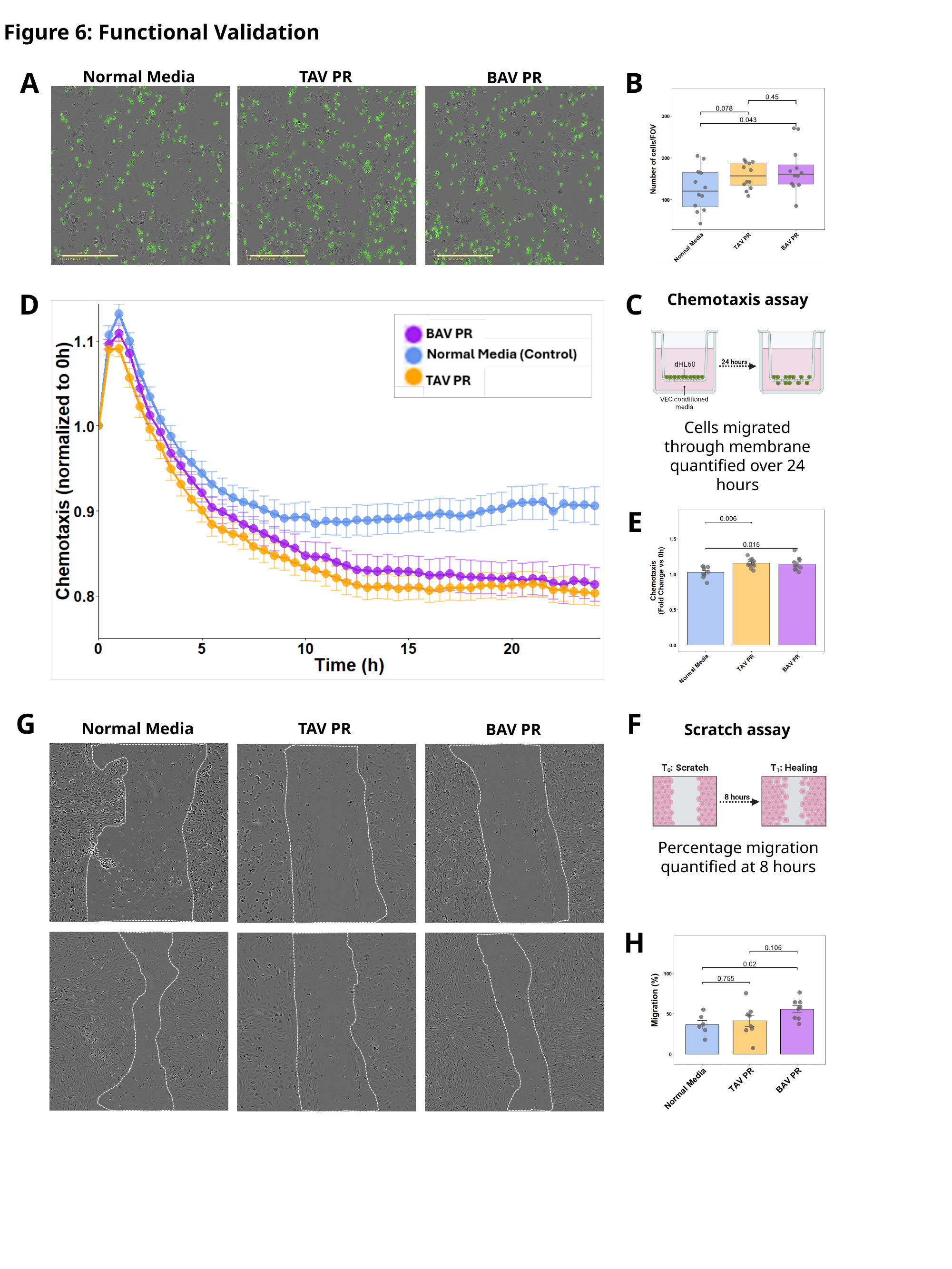

Figure 6: Functional Validation
A
B
Normal Media
TAV PR
BAV PR
D
C
Chemotaxis assay
Cells migrated through membrane quantified over 24 hours
E
G
F
Normal Media
TAV PR
BAV PR
Scratch assay
Percentage migration quantified at 8 hours
H

### Slide 7
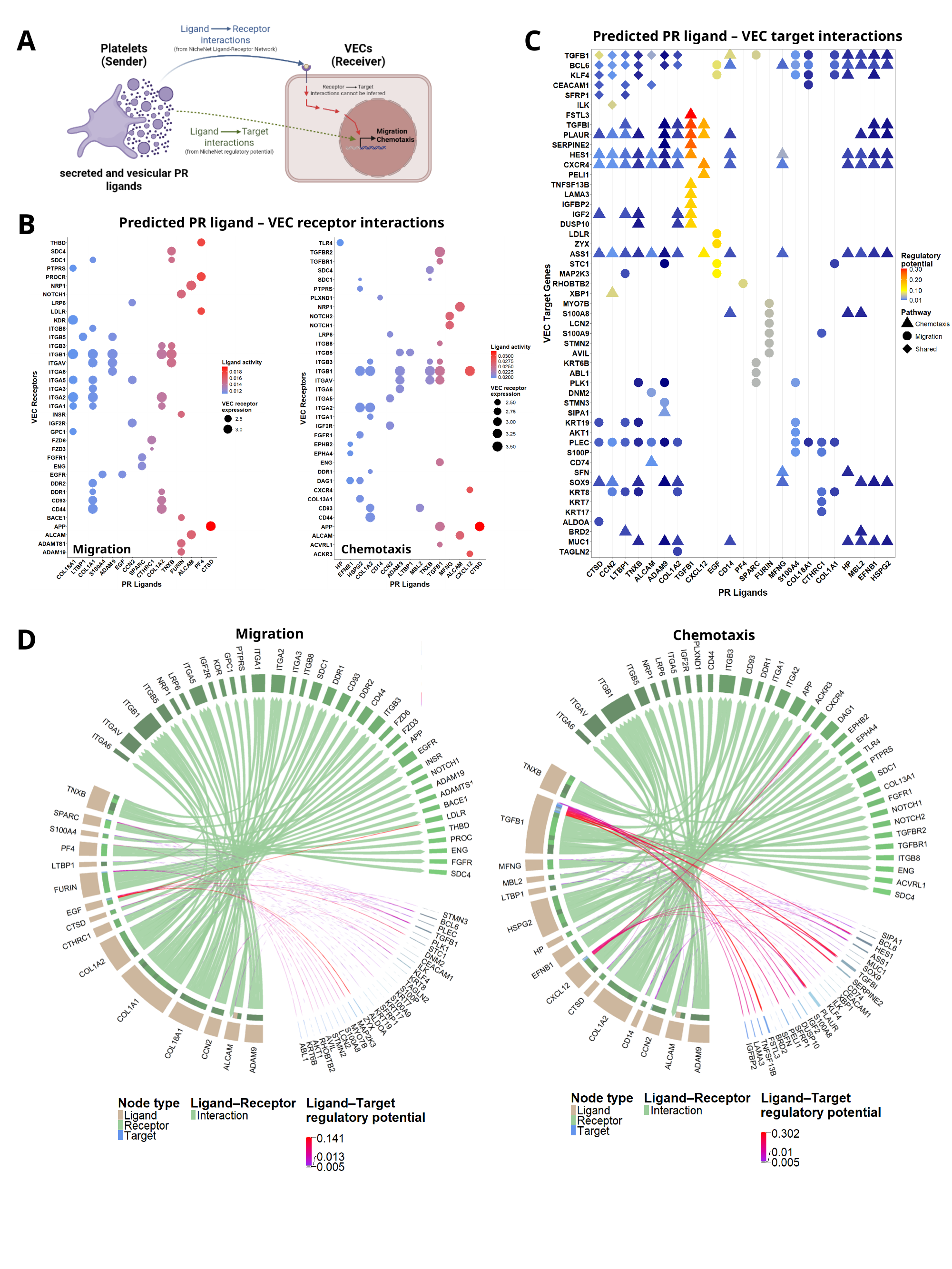

A
C
Predicted PR ligand – VEC target interactions
B
Predicted PR ligand – VEC receptor interactions
Migration
Chemotaxis
D
Migration
Chemotaxis

### Slide 8
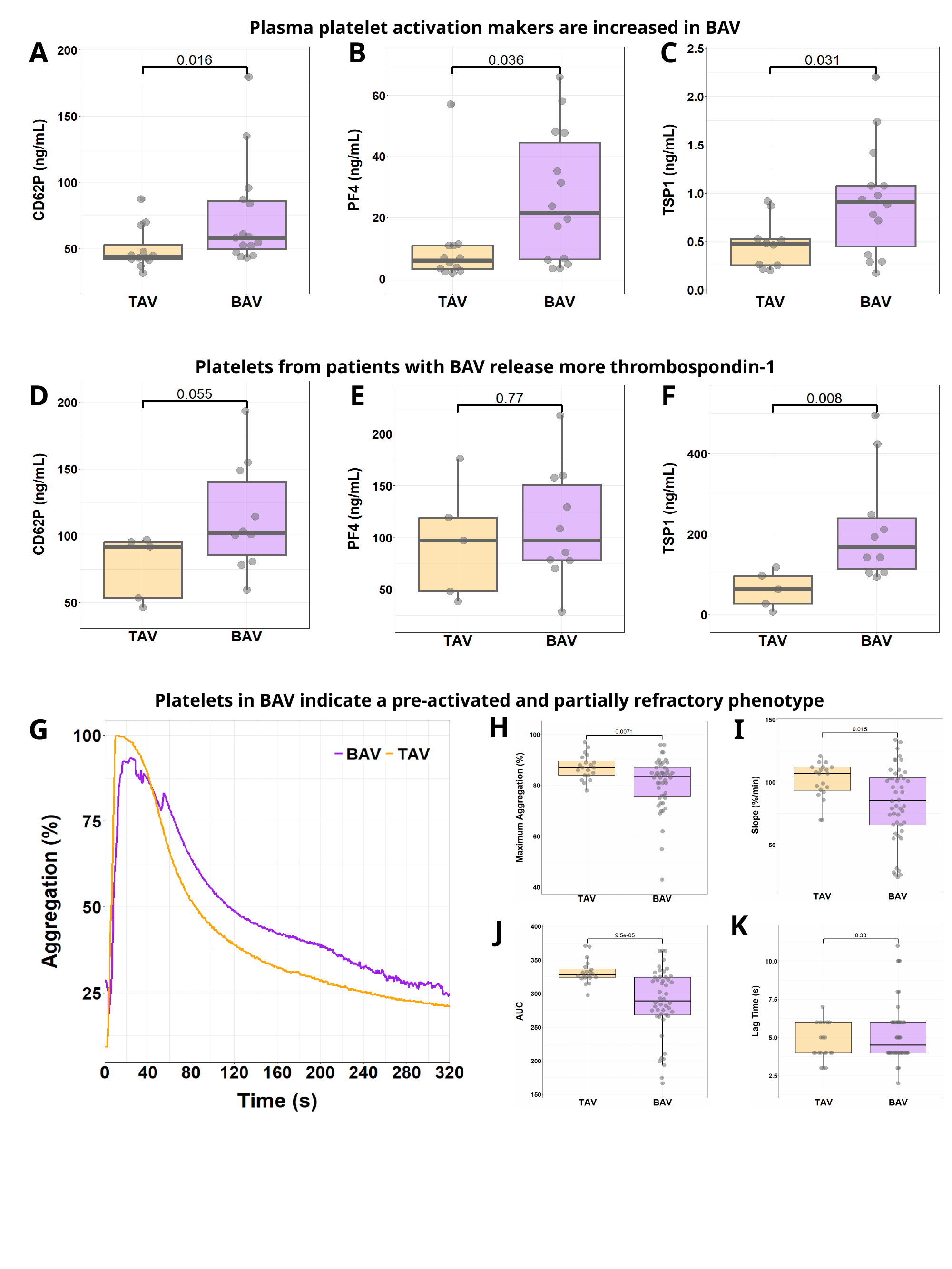

Plasma platelet activation makers are increased in BAV
A
B
C
Platelets from patients with BAV release more thrombospondin-1
D
E
F
Platelets in BAV indicate a pre-activated and partially refractory phenotype
H
I
G
K
J

### Slide 9
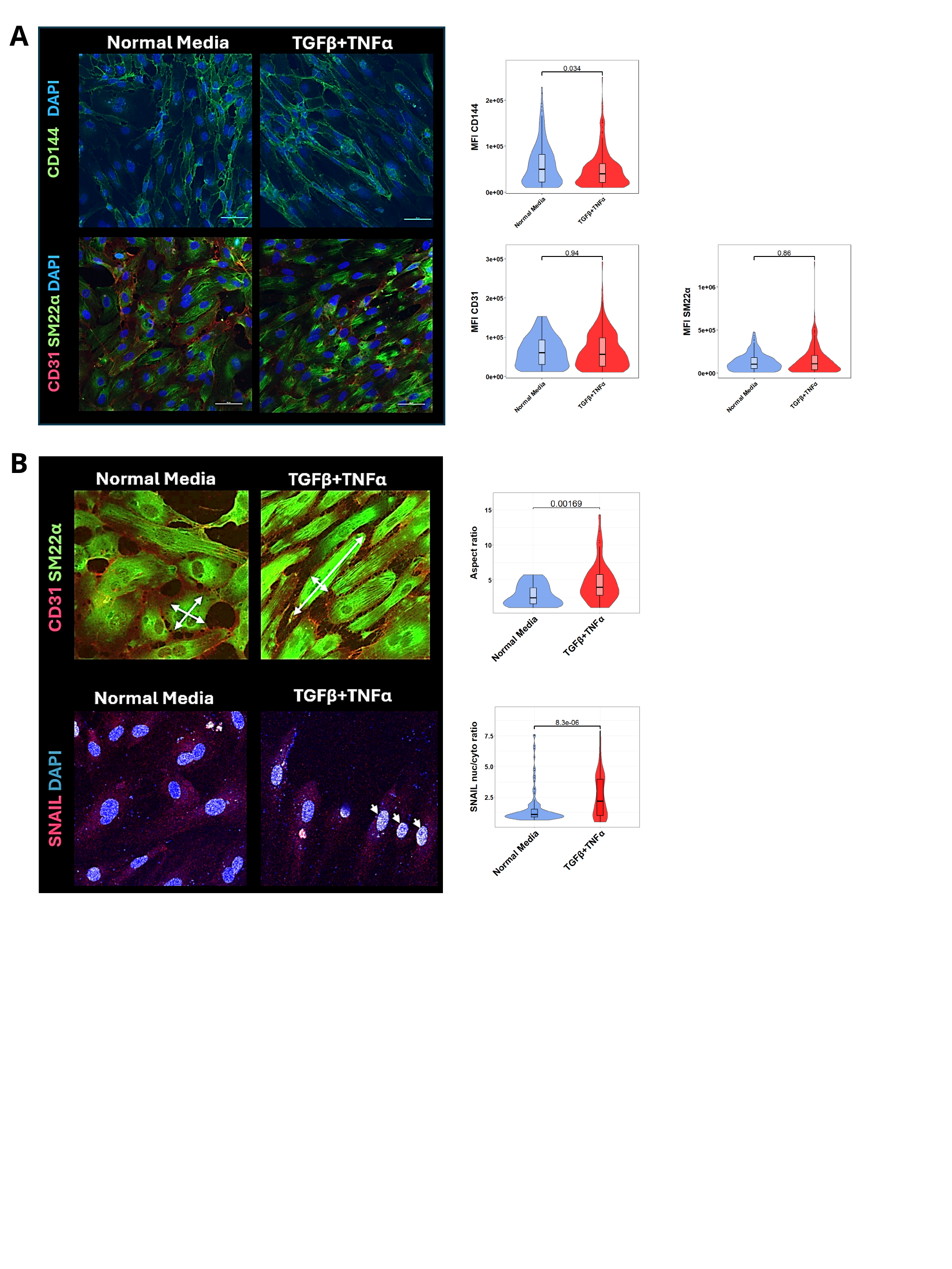

A
B

### Slide 10
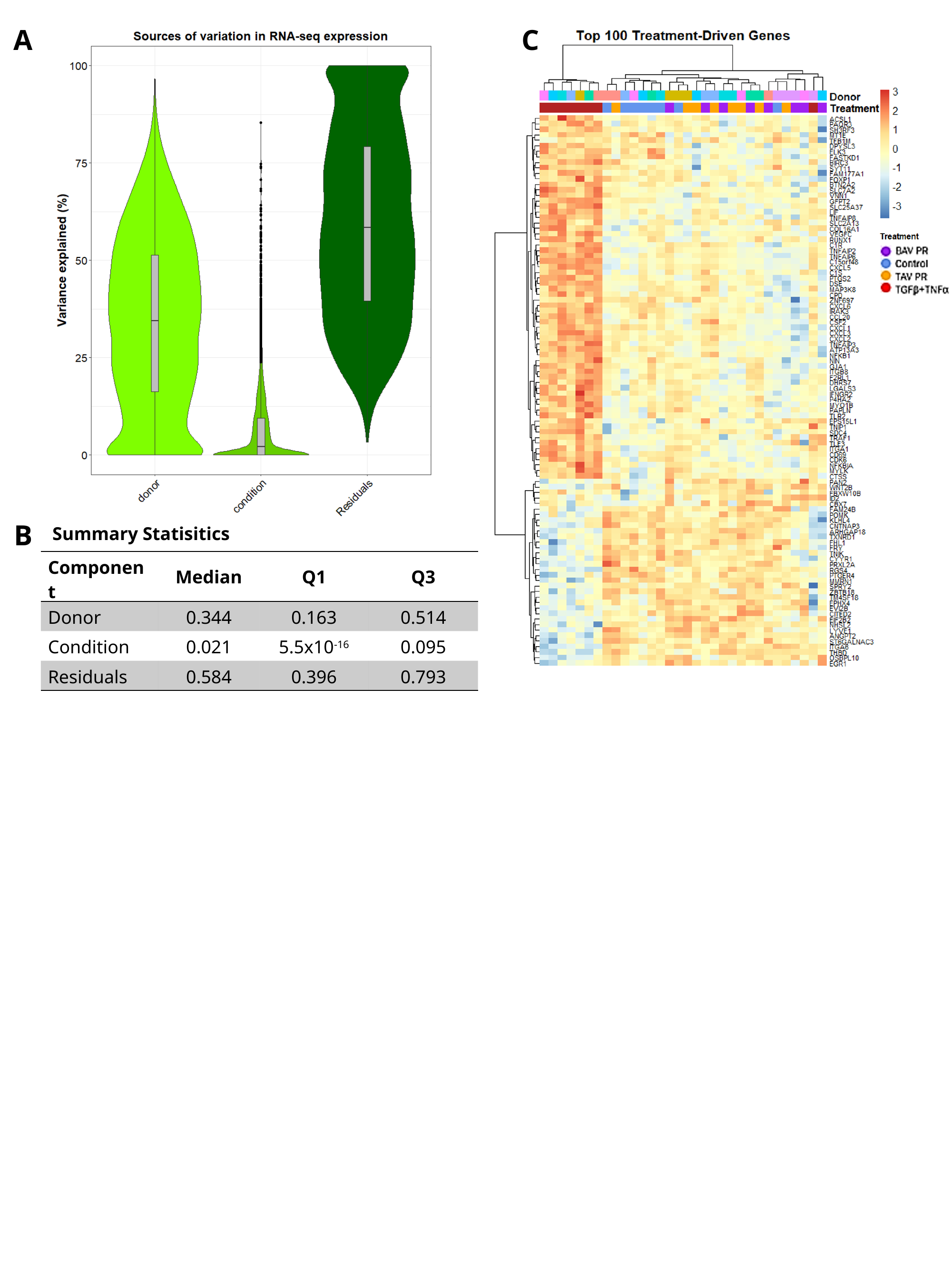

A
C
B
 Summary Statisitics
| Component | Median | Q1 | Q3 |
| --- | --- | --- | --- |
| Donor | 0.344 | 0.163 | 0.514 |
| Condition | 0.021 | 5.5x10^-16^ | 0.095 |
| Residuals | 0.584 | 0.396 | 0.793 |

### Slide 11
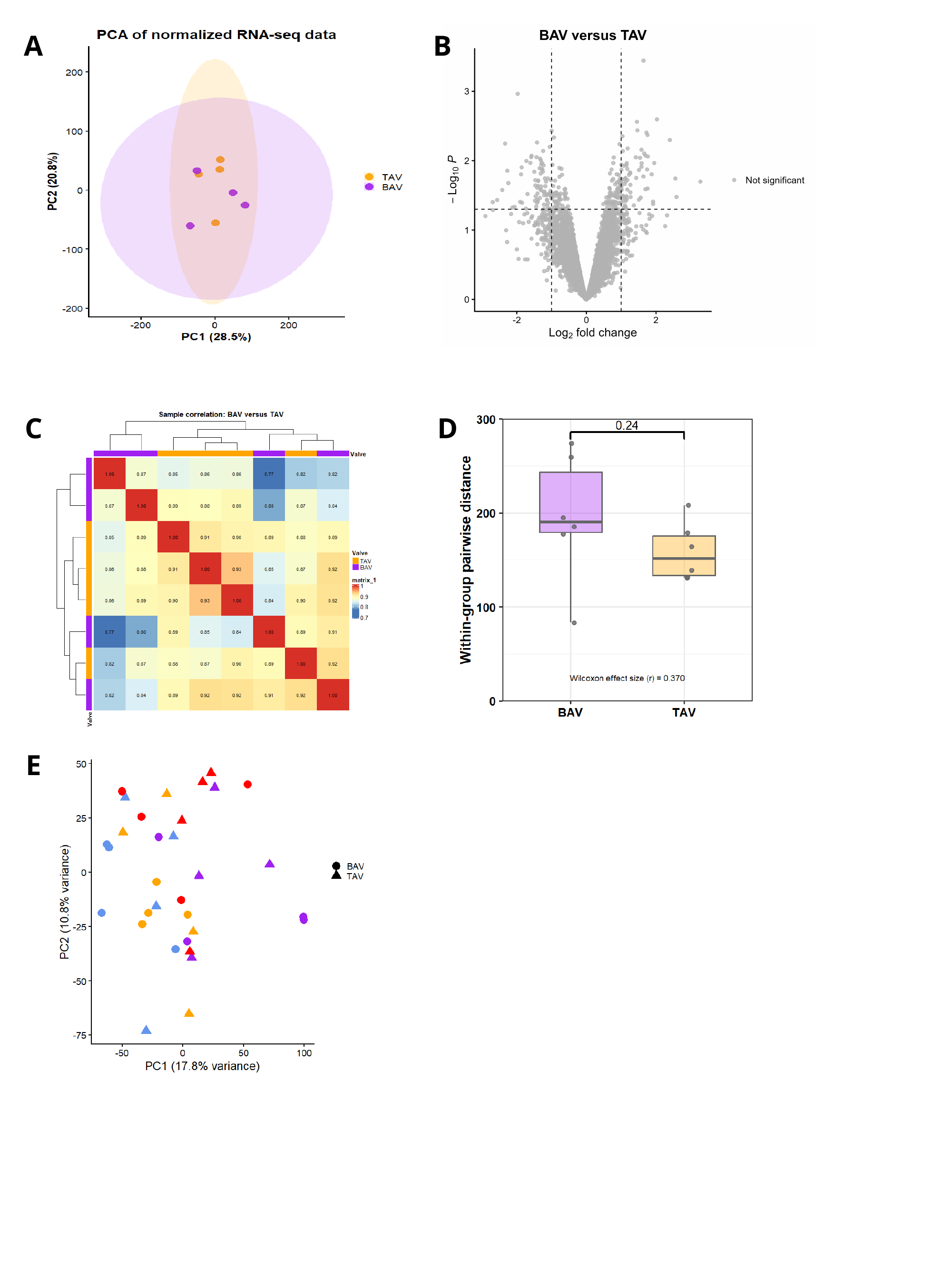

A
B
C
D
E

### Slide 12
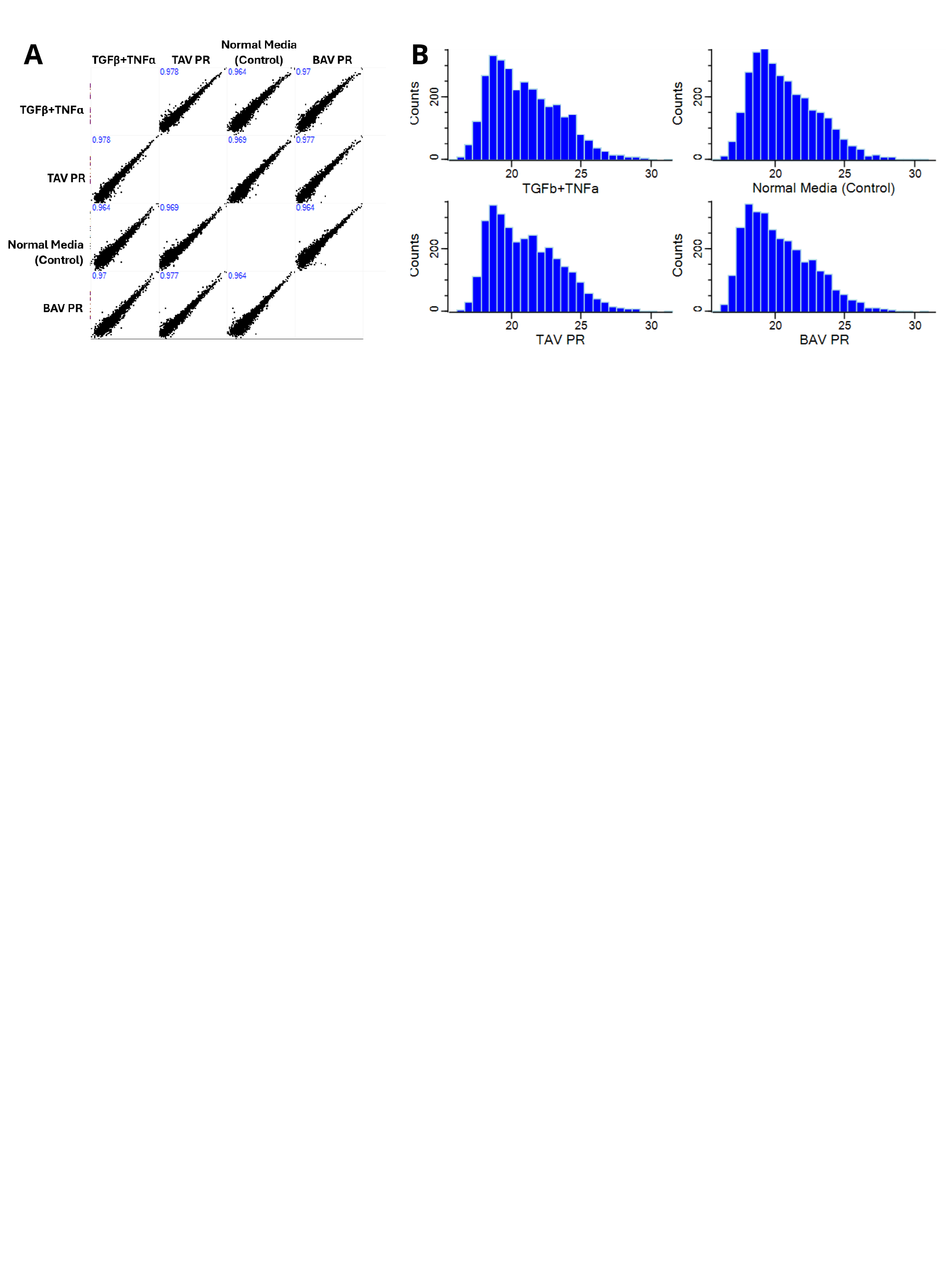

A
B

### Slide 13
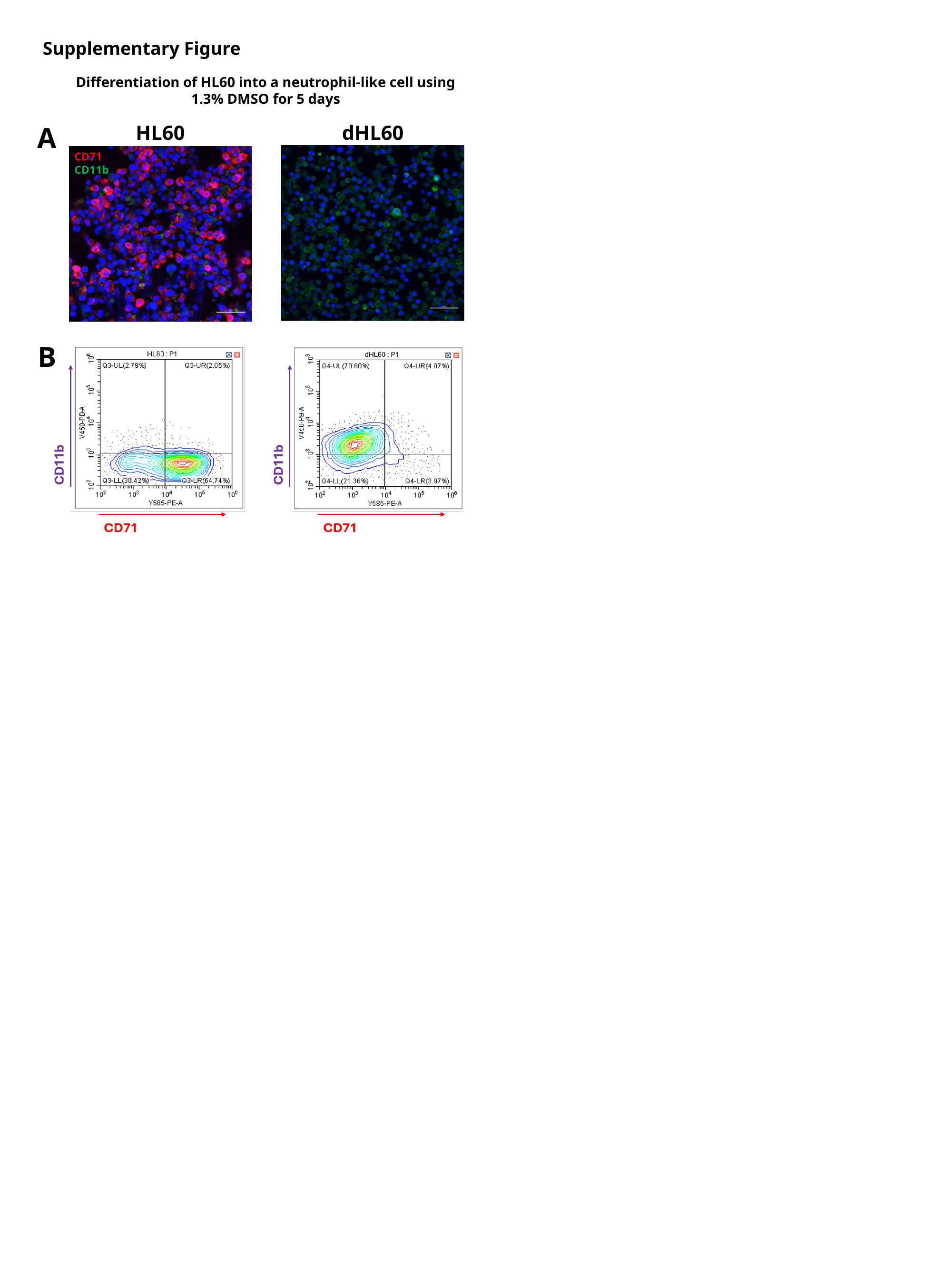

Supplementary Figure
Differentiation of HL60 into a neutrophil-like cell using 1.3% DMSO for 5 days
HL60
dHL60
CD71
CD11b
A
B

### Slide 14
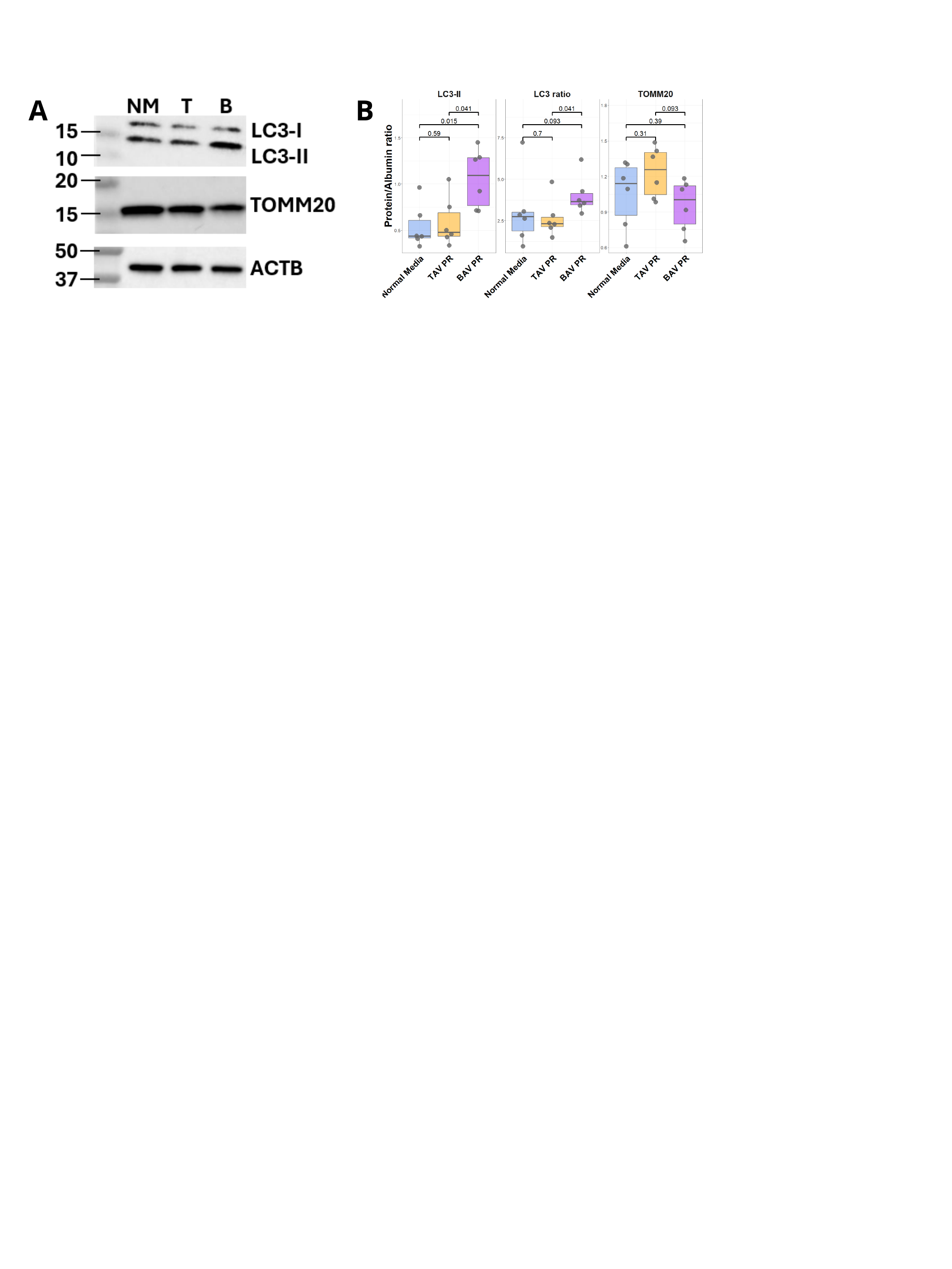

A
B

### Slide 15
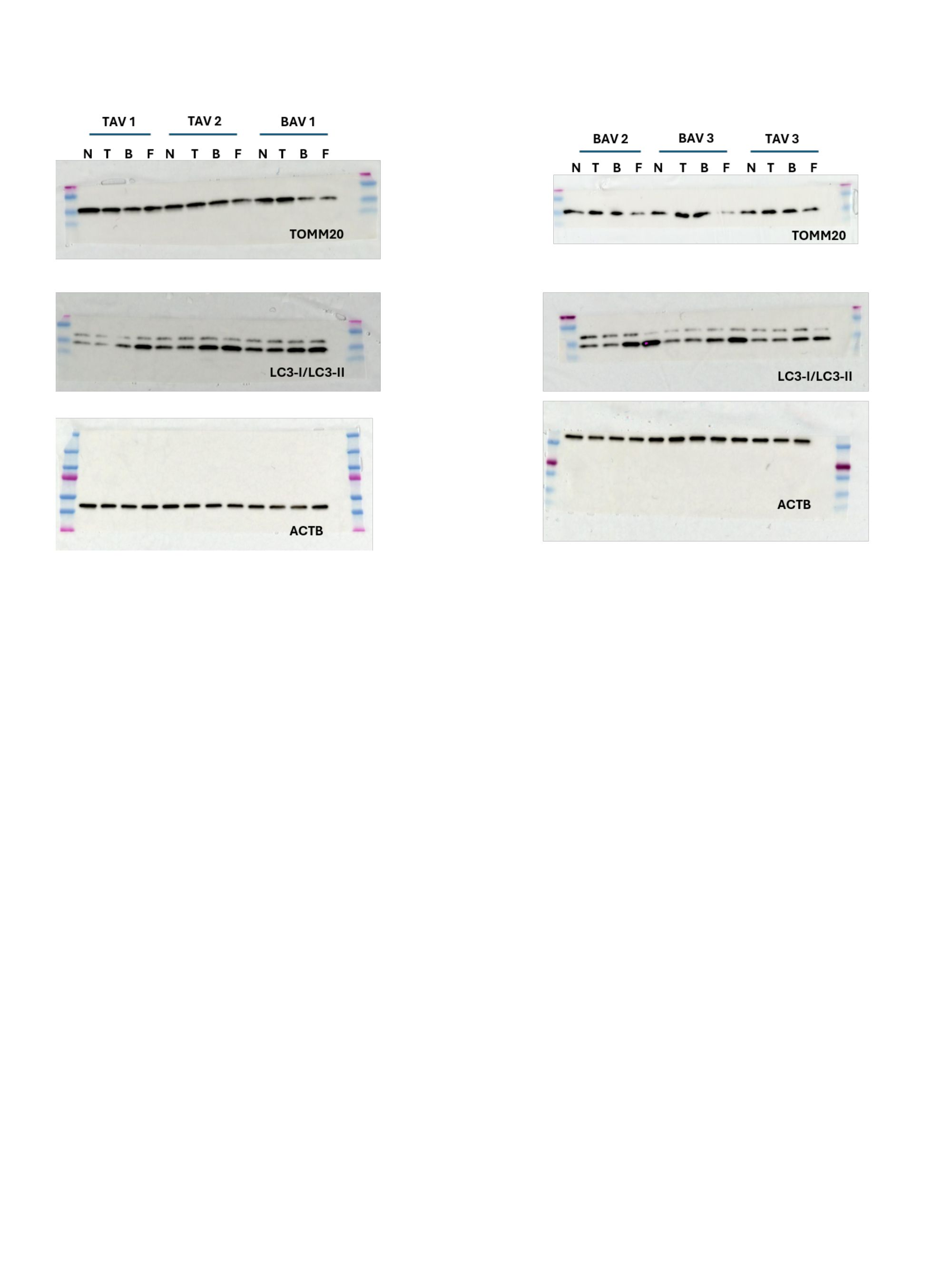
